# A Spatiotemporal Atlas of the Androgen Receptor Proximal Interactome

**DOI:** 10.64898/2026.08.03.742469

**Authors:** Celeste C. Ptak, Conor O’Rourke, Jimmy K. Eng, Lilliana Radoshevich, Michael E. Wright

## Abstract

Androgen receptor-interacting proteins (AR-IPs) number close to 1,000, yet their organization across subcellular space and time remains uncharted. Proximity labeling identifies direct partners and neighboring proteins, thereby expanding AR-IPs to AR-proximal interacting proteins (AR-PIPs). Using proximity labeling quantitative mass spectrometry (PL-qMS), we construct a spatiotemporal atlas of the cytosolic, microsomal, and nuclear compartments in LNCaP prostate tumor cells. PL-qMS recovered 82.2% of the known AR-interactome in extranuclear compartments and 84.2% in the nucleus, identifying 4,751 AR-PIPs that remodel across an androgen time course. The retromer formed an androgen-sensitive AR-proximal interaction network (AR-PIN) verified by proximity ligation assays (PLAs). Moreover, partial VPS26A disruption attenuated androgen-regulated transcription and mislocalized the AR coactivator TMF1, defining a retromer-AR-TMF1 axis. In the nucleus, AR-PINs recover 100% of the Launonen 2021 ChIP-SICAP chromatome and reveal a PLA-verified translation-to-transcription handoff involving eIF4G and 4E-BP1. This spatiotemporal atlas provides a proximal framework for probing AR function in cells.

## Introduction

The androgen receptor (AR) cDNA was cloned nearly four decades ago [1,2], and since then, approximately 1,000 proteins have been identified as AR interactors [3]. AR is a ligand-activated steroid hormone transcription factor that governs male development, reproductive physiology, and prostate cancer progression in men [4]. Moreover, recent studies have shown an increasing role for AR signaling in non-reproductive organs and tissues [5]. AR interactors, better known as AR-interacting proteins (AR-IPs), are thought to regulate the specificity of AR signaling, and their cell-, tissue-, and organ-specific expression is a critical determinant of AR signaling outcomes. AR-IPs span diverse functional classes, including coregulators, chaperones, kinases, pioneer factors, and trafficking factors, identified through successive waves of methodology, from yeast two-hybrid assays, recombinant GST fusions, and phage display [6,7,8,9] to antibody-directed co-immunoprecipitation and affinity-purification mass spectrometry platforms such as RIME [10,11], and most recently to proximity labeling [12]. These efforts have produced a curated AR-IP library that now encompasses nearly 1,000 proteins [3]. Yet the biologically active AR-interactome is dynamic and context-specific, governed by AR-IP expression across tissues, developmental contexts, and disease states [5,4], and the catalog itself offers limited predictive power. Knowing that a protein binds AR does not reveal where in the cell the interaction occurs, when during the signaling cascade it assembles, or whether it functionally contributes to AR-dependent transcription.

AR is a ligand-activated hub protein that orchestrates transcriptional programs at more than 10,000 androgen response elements across the human genome [13,14], with most androgen-regulated genes depending on the assembly of coregulators, chromatin remodelers, Mediator subunits, and pioneer factors at distal enhancers [15,16]. Beyond this canonical chromatin axis, an expanded view of AR action has emerged, extending into non-coding RNA biology [17,18,19], alternative splicing, alternative polyadenylation, and translational control [20,21,22,23,24]. This expanded repertoire implies that the functionally relevant AR-interactome reaches well beyond chromatin-bound coregulators. The receptor itself is built for such breadth, with an intrinsically disordered N-terminal domain that mediates multivalent interactions that drive AR condensate formation and optimal transcriptional activity [25,26,27,28]. Upon ligand binding, AR then undergoes one-way cytosolic-to-nuclear trafficking [29,30], providing ample opportunity to engage diverse proteins and complexes en route to chromatin. Together, these properties raise a central question of when and where each class of partner physically engages AR within the cell.

Proximity labeling captures both direct and neighboring interactions within approximately 200 nm of a target receptor [12,31,32,33,34], expanding the observable interactome beyond what binary methods alone capture. Coupled to quantitative mass spectrometry and data-independent acquisition, proximity labeling routinely quantifies thousands of proteins per run with high reproducibility [35,36,37,38]. The resulting proximal proteome includes known, presumably functional interactors, as well as neighboring proteins that reflect a receptor’s subcellular localization, trafficking, metabolism, and degradation [39]. Rather than treating these neighbors as uninformative bystanders, we propose that the full proximal population constitutes a form of subcellular cartography, an expression-dependent landscape in which spatial co-residence and low-affinity interactions [40,41] among neighboring proteins shape signaling at the target receptor. A bystander is not noise; a protein positioned within the AR proximal environment carries the latent potential to modulate AR signaling should its expression or localization become dysregulated, as frequently occurs in prostate cancer.

Here, we leverage proximity labeling quantitative mass spectrometry (PL-qMS) [42,43] to construct a spatiotemporal atlas of the AR proximal interactome, resolved over a minute-scale androgen stimulation time course, across the cytosolic, microsomal, and nuclear compartments of human prostate tumor cells. We define the proteins that populate the AR-proximal environment as AR-proximal interacting proteins (AR-PIPs), encompassing known AR-IPs and previously uncharacterized proximal proteins alike, and we define their assembly into functional complexes as AR-proximal interaction networks (AR-PINs). PL-qMS recovers the majority of the known AR-interactome in every compartment and resolves the order in which coregulator classes, chromatin remodelers, Mediator subunits, and transcription- and translation-coupled factors enter and exit the proximal interactome. Within this atlas, we identify and functionally validate two AR-proximal programs. In the extranuclear compartments, we uncover a novel proximal interaction between AR and the retromer complex. Retromer integrity, through VPS26A, is required for optimal AR-dependent transcription of canonical androgen-regulated genes that correlates with Golgi localization of the AR coactivator TMF1. In the nucleus, we resolve an early, transient engagement of AR with the cap-binding translational machinery and validate it by proximity ligation. Cross-validation of nuclear AR-PIPs against an independent ChIP-SICAP chromatome recovers all previously validated chromatin-bound AR interactors [44]. Together, the atlas and these two validated networks position subcellular proximal proteomes as an entry point for probing AR function, shifting AR biology from cataloging individual interactions toward mapping the proximal networks that govern androgen-dependent signaling and its dysregulation in disease.

## Results

### A spatiotemporal subcellular atlas of the AR proximal interactome

AR’s primary function is to act as a ligand-activated nuclear transcription factor [4], but years of biochemical research have shown that AR also engages numerous extranuclear proteins across cytosolic and membrane compartments [45,46,47,48]. Many of these extranuclear interactions are coregulators of AR-mediated transcription, which remain poorly annotated to date. To comprehensively elucidate both direct and indirect proximal AR interactors in the extranuclear and nuclear spaces as AR responds to androgen, a stable LNCaP cell line derivative called LNCaP-APEX2-AR cells was developed that expresses an APEX2-AR fusion protein under doxycycline-inducible control (Figures 1A and 1B)[49,50]. We selected the APEX2 peroxidase to capture proximal interactors, generating covalent biotin labels on proteins within ∼200 nm of the APEX2-AR fusion under both acute and chronic androgen exposure. APEX2 was appended to the N-terminus of full-length AR, retaining all functional domains (i.e., NTD, DBD, hinge, and LBD) (Figure 1A). Unlike constitutively active TurboID, APEX2 biotinylation requires acute exposure to hydrogen peroxide, enabling capture of spatiotemporally proximal interactions with both unliganded and liganded AR.

**Figure 1.**
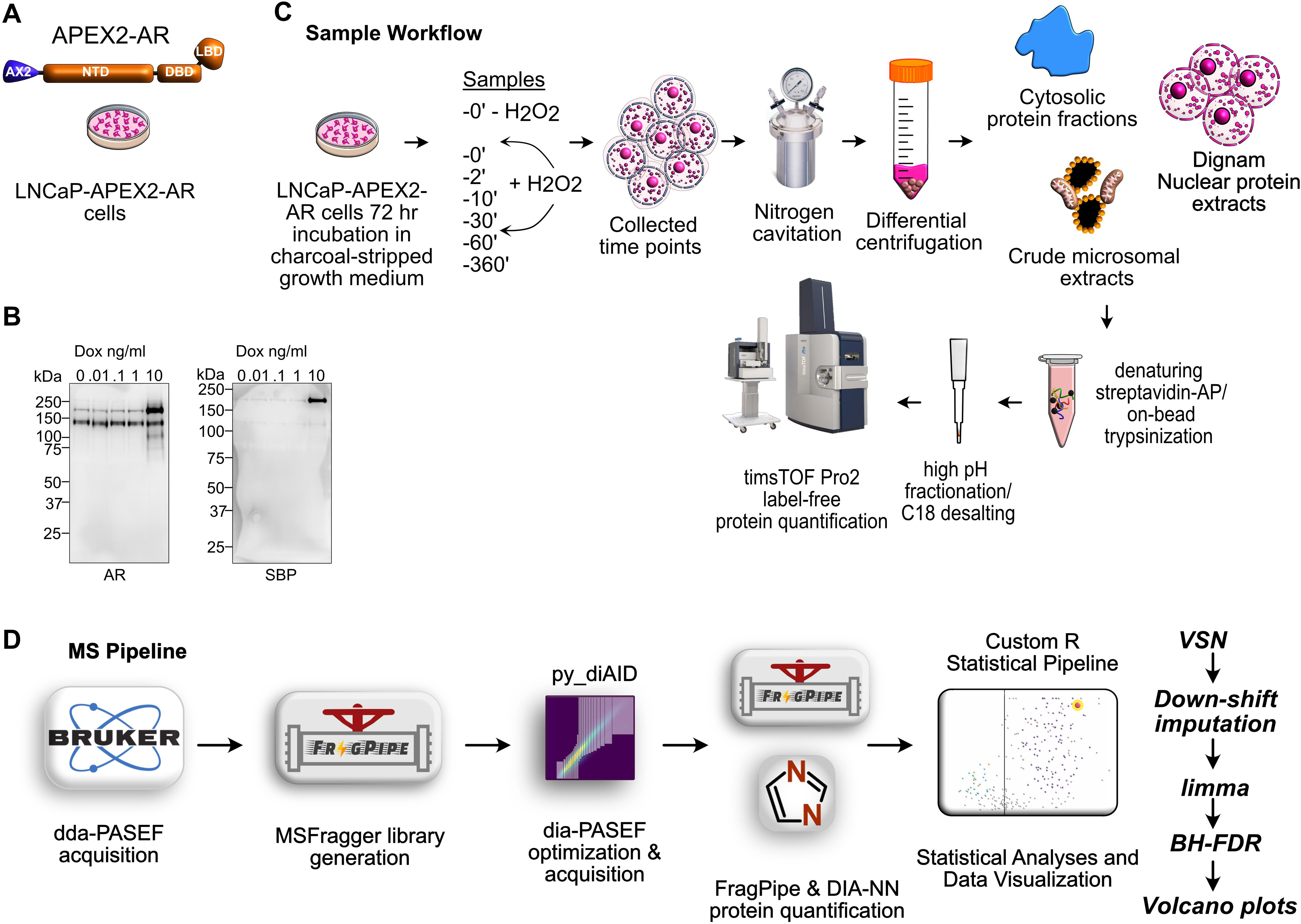
Defining the proximal AR-interactome by proximity labeling quantitative mass spectrometry (PL-qMS) (A) Molecular structure of APEX2-AR, in which APEX2 is fused to full-length AR retaining the NTD, DBD, hinge, and LBD, expressed in LNCaP-APEX2-AR prostate tumor cells. (B) Doxycycline dose-dependent induction of APEX2-AR detected by anti-AR and anti-SBP western blot (0.01, 0.1, 1, and 10 ng/ml doxycycline). (C) Sample workflow. LNCaP-APEX2-AR cells were propagated for 72 hr in charcoal-stripped growth medium; a minus-H2O2 control and H2O2-pulsed time points (0, 2, 10, 30, 60, and 360 min) were collected and processed by nitrogen cavitation and differential centrifugation into cytosolic, crude microsomal, and Dignam nuclear extracts, followed by denaturing streptavidin-affinity purification with on-bead trypsinization, high-pH fractionation and C18 desalting, and timsTOF Pro2 label-free protein quantification. (D) MS pipeline. dda-PASEF acquisition, MSFragger spectral-library generation, py_diAID dia-PASEF optimization and acquisition, FragPipe and DIA-NN protein quantification, and a custom R statistical pipeline (VSN normalization, down-shift imputation, limma, and BH-FDR) that produced the volcano plots. See also Supplementary Fig. 1-3.

For the inducible APEX2-AR fusion to faithfully report AR biology, it must behave like endogenous AR and undergo cytoplasmic-to-nuclear trafficking in response to androgen. We first tested APEX2-AR trafficking behavior using immunofluorescence microscopy, staining for AR, the SBP tag, and biotin, and verified that it behaved like endogenous AR. APEX2-AR was cytoplasmic in androgen-depleted cells and underwent androgen-mediated cytoplasmic-to-nuclear translocation upon androgen stimulation, with the biotin signal tracking APEX2-AR from the cytosol into the nucleus over the time course (Supplementary Fig. 1). Next, APEX2-AR trafficking was corroborated biochemically by Western blot analysis of subcellular fractionated protein extracts. Across the androgen time course, APEX2-AR progressively redistributed from the cytosolic and membrane compartments (Supplementary Fig. 2). Robust biotinylation activity was also detected in biotinylated substrate proteins across all compartments throughout the androgen time course (Supplementary Fig. 2). Together, the immunofluorescence and fractionation results establish that the inducible APEX2-AR system replicates the ligand-dependent cytoplasmic-nuclear trafficking of endogenous AR, validating it as a faithful experimental system for capturing AR proximal interactors under both acute and chronic androgen stimulation.

To profile AR proximal interactors across the androgen response, LNCaP-APEX2-AR cells were grown in androgen-depleted growth medium (*i.e.*, 72 hrs), treated with doxycycline (*i.e.*, 24 hrs), incubated with biotin-phenol (*i.e.*, 1 hr), and exposed to synthetic androgen (*i.e.*, 100 nM R1881) under multiple time points (*i.e.*, 0, 2, 10, 30, 60, and 360 minutes). This strategy would capture unliganded extranuclear interactors, as well as chronically liganded nuclear proximal interactors of APEX2-AR. Cells were fractionated by nitrogen cavitation and differential centrifugation into cytosolic, crude microsomes (*i.e.*, membrane fraction), and nuclear fractions (Figure 1C).

We developed a proteomic workflow to quantify all proximity-labeled proteins by label-free quantitative mass spectrometry on the timsTOF Pro2 platform (Figure 1C). Biotinylated proteins from each fraction were affinity-purified on magnetic streptavidin beads under denaturing conditions, and streptavidin immunoblotting confirmed specific enrichment of biotinylated proteins across the time course (Supplementary Fig. 3). Since streptavidin-biotin interactions are nearly irreversible, the magnetic beads were subjected to single-pot solid-phase-enhanced sample preparation (SP3) under stringent wash conditions (*e.g.*, 1% SDS, 4 M Urea), followed by on-bead trypsin digestion and high-pH peptide fractionation prior to LC-MS/MS. To maximize proteome depth and quantitative accuracy, we employed a two-pass mass spectrometry pipeline (Figure 1D). Each biological replicate was interrogated by data-dependent acquisition parallel-accumulation-serial-fragmentation (dda-PASEF) to generate comprehensive compartment-specific spectral libraries via MSFragger [51], which were then queried by data-independent acquisition (dia-PASEF) on all three biological replicates using py_diAID-optimized ion selection [52]. Protein identification and quantification were performed with FragPipe and DIA-NN [53,54,55,56], and quantifications were analyzed with a single custom R pipeline applied identically to all three compartments (i.e., variance-stabilizing normalization, down-shift imputation, limma moderated t-test, and Benjamini-Hochberg FDR) (Figure 1D).

Volcano plots of three replicate PL-qMS experiments revealed robust enrichment of streptavidin-purified proteins in all three compartments across all hydrogen-peroxide-treated time points (*i.e.*, 0 through 360 minutes) relative to the background control (i.e., 0 minutes, no H_2_O_2_) (Figure 2). In every experimental sample relative to the background control, APEX2-AR was among the most significantly enriched proteins, confirming the specificity and sensitivity of the proximity-labeling workflow. Applying a threshold of an adjusted p-value of at most 0.05 and a log2 fold change greater than 1 to classify proteins as significantly enriched at each time point, the entire population of statistically enriched proteins was classified as “AR proximal interacting proteins” (AR-PIPs). This AR-PIP population encompassed the curated 989-member AR-interactome cataloged in BioGRID and other protein databases [3] (Supplementary Data 4). Known members of the AR-interactome were consistently enriched in the cytosolic (Supplementary Data 1), microsomal (Supplementary Data 2), and nuclear (Supplementary Data 3) fractions and were broadly reproducible across time points, indicating stable, reproducible labeling over the 360-minute time course. In total, we identified 2,994 cytosolic, 3,475 microsomal, and 3,378 nuclear AR-PIPs, which together define a union of 4,751 proteins proximal to AR. Unequivocally, these subproteomes represent the most comprehensive spatiotemporal map of the AR proximal interactome assembled to date.

**Figure 2.**
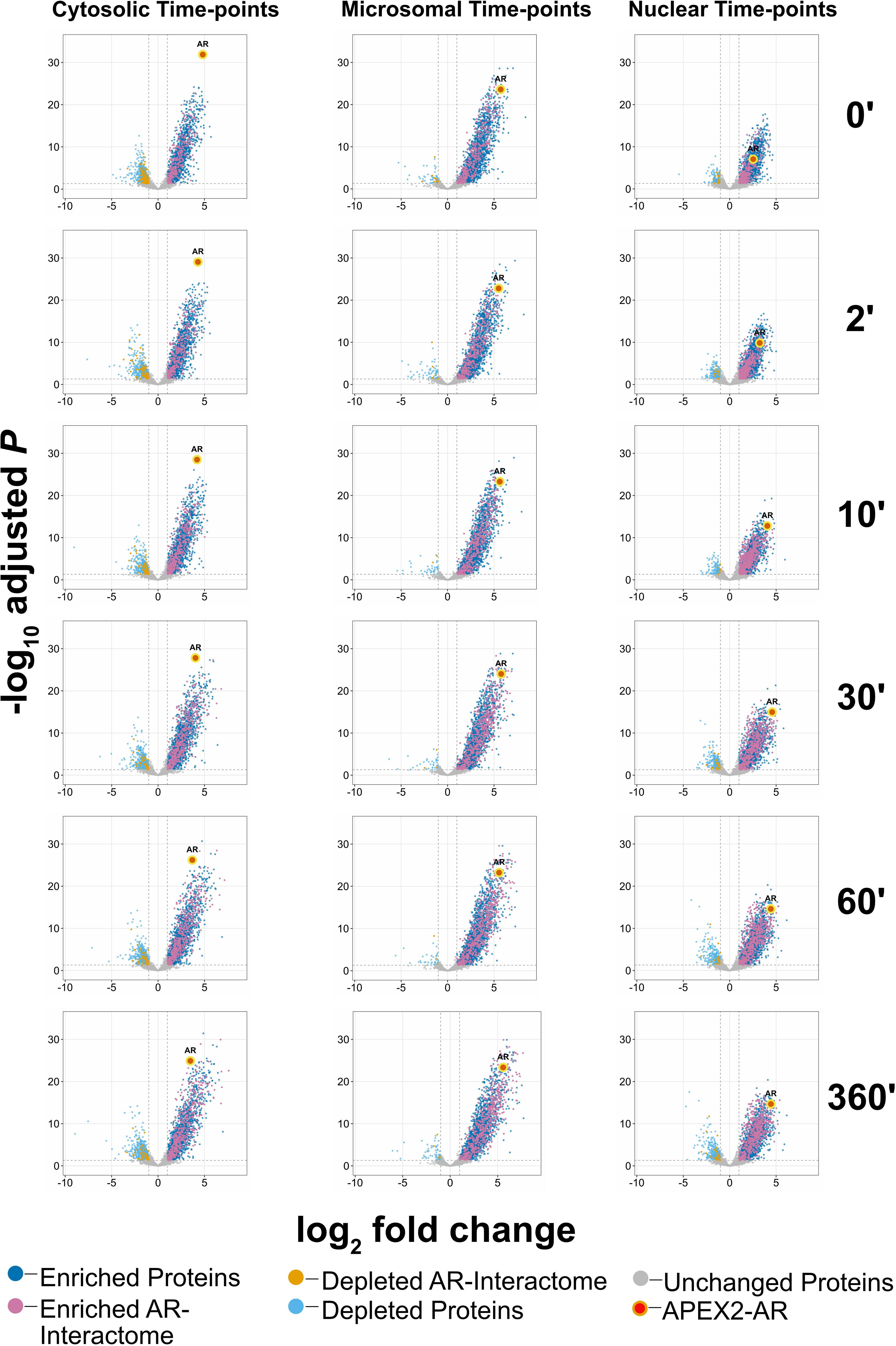
Volcano plots of quantified cytosolic, microsomal, and nuclear proximal proteomes across the androgen time course. Time-course volcano plots of proteins quantified by PL-qMS, arranged as cytosolic (left), microsomal (center), and nuclear (right) fractions in columns at 0, 2, 10, 30, 60, and 360 min of 100 nM R1881 androgen treatment in rows. For each fraction, the minus-H2O2 background control (0 min, no hydrogen peroxide) was used to quantify enrichment across the hydrogen peroxide-treated time points; three biological replicates per condition. Axes: log2 fold change (x) versus −log10 adjusted P (y). Threshold, adjusted P ≤ 0.05 and log2 fold-change > 1. Circles: dark blue, enriched proteins; pink, enriched AR-interactome proteins; orange, depleted AR-interactome proteins; light blue, depleted proteins; gray, unchanged proteins; red-outlined circle, APEX2-AR. Source data, Supplementary Data 1-3 and the Source Data file.

### The AR-interactome is recoverable and enriched subproteome across all three compartments

A proximity atlas is only as informative as its ability to recover known biology, so we first tested how completely and how specifically the workflow captured the curated 989-member AR-interactome. To formally test whether known AR-interactome proteins were enriched among the biotinylated proximal proteins, we applied one-sided hypergeometric tests across two increasingly stringent comparisons. First, the overall detection of the AR-interactome was evaluated across all proteins quantified by PL-qMS to assess the sensitivity of our approach in verifying the proximity of known AR-interactome members. Second, we determined per-time-point over-representation of the AR-interactome among the significantly enriched AR-PIPs, because known members of the AR-interactome are expected to be proximal and thus should be enriched after affinity purification. Evaluated against the LNCaP-expressed universe of approximately 12,000 proteins, the nuclear PL-qMS workflow detected and recovered 833 of 989 AR-interactome members (84.2% coverage; p = 1.9 × 10⁻¹¹⁵), and the extranuclear workflow detected 813 of 989 (82.2%), of which 666 (81.9%) were verified as significantly enriched AR-PIPs in one or both extranuclear compartments (Figure 3A to 3C).

**Figure 3.**
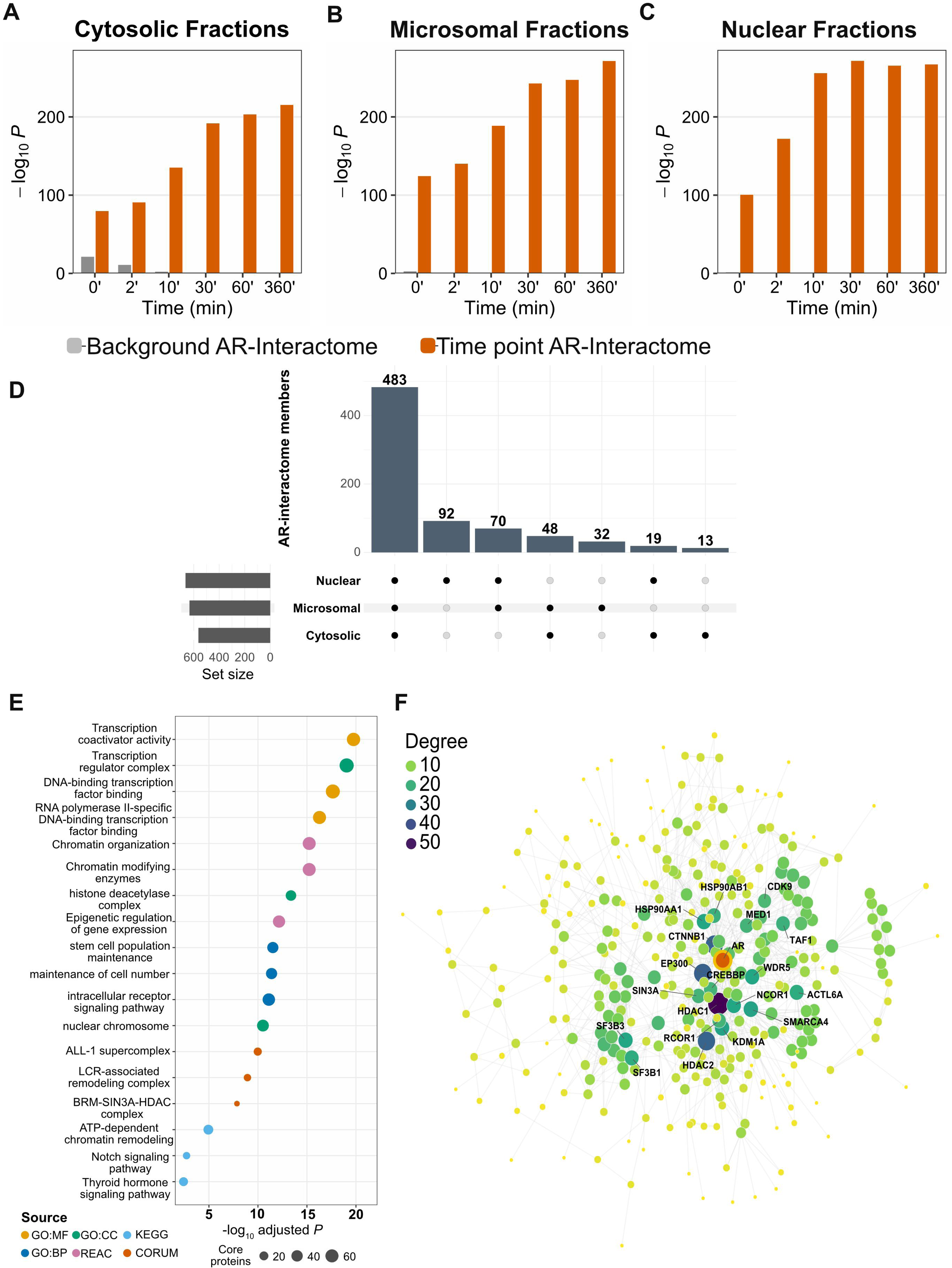
The AR-interactome is a recoverable and enriched subproteome across all three compartments. (A-C) Hypergeometric test for enrichment of the 989-member AR-interactome reference within the proximal AR-PIP set (adjusted P ≤ 0.05 AND log2 fold-change > 1) at each time point, plotted as the −log10 P-value, for the (A) cytosolic, (B) microsomal, and (C) nuclear fractions. Orange bars (Time point AR-Interactome) show enrichment in the hydrogen peroxide-treated, androgen-stimulated samples; gray bars (Background AR-Interactome) show negative-control samples lacking hydrogen peroxide, confirming that the AR-interactome signal is APEX2-dependent. (D) UpSet plot of the 989-member AR-interactome across the cytosolic, microsomal, and nuclear compartments; 483 members are shared by all three compartments. (E) g:Profiler functional enrichment of the recovered AR-interactome, plotted as −log10 adjusted P; points colored by source (GO:MF, GO:CC, GO:BP, KEGG, REAC, CORUM) and sized by the number of core proteins. (F) Cytoscape network of the recovered core AR-interactome, with nodes colored by degree and sized by the number of core proteins; hub coregulators (e.g., AR, CREBBP, EP300, HDAC1, NCOR1, MED1) are labeled. Source data in Supplementary Data 4, 5, 6, 7, and 8, and the Source Data file.

Beyond detection, enrichment of the AR-interactome was decisive. For example, AR-interactome members among the nuclear AR-PIPs rose progressively within each time point, from 17.1% at 0 minutes (334 of 1,958, p = 1.9 × 10⁻¹⁰¹) through 22.9% at 30 minutes (571 of 2,490, p = 9.5 × 10⁻²⁷³) to 23.8% at 360 minutes (550 of 2,315, p = 2.9 × 10⁻²⁶⁸). Similarly, the cytosolic and microsomal fractions showed the same highly significant over-representation across the time course (Figure 3A-3C). This progressive enrichment is decisive evidence that the atlas concentrates the known AR-interactome above background at every stage of the androgen response, because random contamination would neither enrich a curated interactome nor show monotonically increasing significance.

To identify AR-interactome members that AR engages regardless of subcellular location, we integrated data from the three compartments into a single analysis. Of the 989 curated members, 757 were significantly enriched as AR-PIPs in at least one compartment, and 483 were enriched in all three, defining a compartment-independent core of the AR proximal interactome (Figure 3D)[57].

This finding prompted us to test whether this shared core reconstructs canonical AR biology rather than a generic abundant-protein background. Thus, we characterized the 483 all-compartment core by functional enrichment and network connectivity. g:Profiler analysis recovered the defining machinery of AR-driven transcription, including transcription coactivator activity, the transcription regulator complex, DNA-binding transcription factor binding, chromatin organization and chromatin-modifying enzymes, and the histone deacetylase complex (Figure 3E)[58]. A STRING physical interaction network of the core was densely interconnected and centered on AR, with the highest-degree hubs comprising established AR coregulators, including CREBBP, EP300, HDAC1, HDAC2, SIN3A, NCOR1, KDM1A, SMARCA4, and MED1 (Figure 3F)[59]. The atlas therefore not only detects the known AR-interactome but is enriched in every compartment and reconstructs its canonical transcriptional-coregulator architecture, establishing a validated foundation for the compartment-specific analyses that follow.

### Temporal remodeling of the extranuclear AR proximal interactome

Having established the scope and specificity of the atlas, we asked whether the extranuclear AR proximal interactome was remodeled as AR traffics from the cytoplasm into the nucleus. We first quantified when each protein enters the interactome by assigning it to the earliest time point at which it becomes enriched as a proximal interactor. Most extranuclear AR-PIPs were already proximal to AR at the unliganded baseline, and successive waves of proteins first became proximal at later time points, with the largest cohort of new entrants appearing at 30 minutes (Figure 4A). To ask how the architecture of the interactome itself changed, we constructed STRING physical interaction networks of the shared extranuclear proximal interactors at each time point[59] and tracked their sizes relative to the unliganded baseline. The number of nodes increased only modestly, but the number of edges rose sharply to an approximately three-fold peak at 30 minutes before relaxing (Figure 4B), indicating a transient burst of interconnectivity within the extranuclear interactome that coincides with the peak of new protein entry.

**Figure 4.**
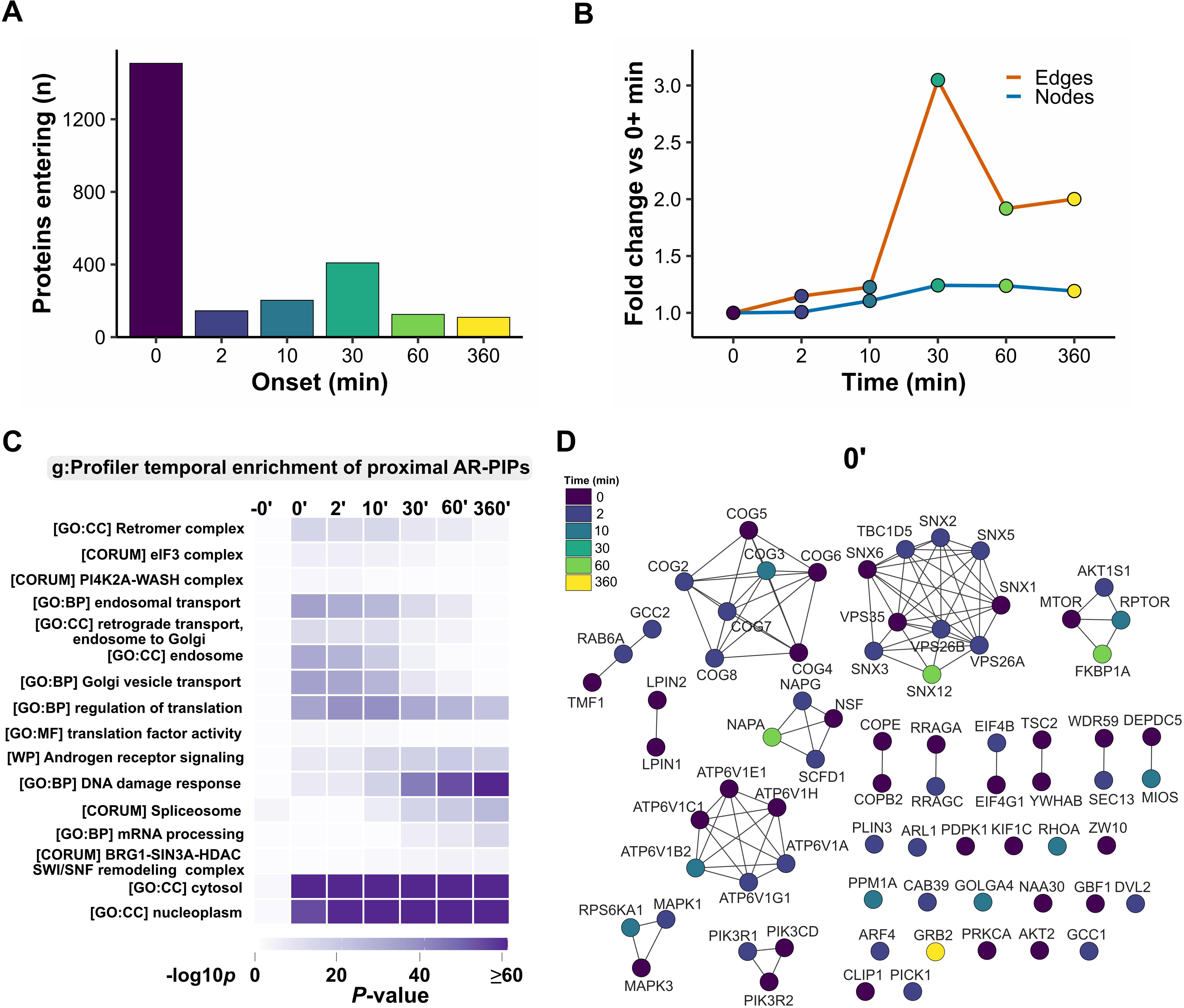
Temporal remodeling of the extranuclear AR proximal interactome. (A) Number of proteins entering the extranuclear proximal interactome by onset time point (0, 2, 10, 30, 60, and 360 min), with the largest cohort entering at 0 min and a second wave at 30 min. (B) Fold change in network edges (orange) and nodes (blue) relative to 0+ min across the androgen time course, with edge density peaking at 30 min. (C) g:Profiler temporal enrichment heatmap of the proximal extranuclear AR-PIPs (rows, selected GO, CORUM, Reactome, and WikiPathways terms including the retromer complex, endosomal and retrograde transport, regulation of translation, and DNA damage response; columns, time points), colored by −log10 P. (D) STRING physical-interaction network of the extranuclear AR-PIPs (representative 0-min layout) visualized in Cytoscape, with clusters for the retromer and sorting nexins, COG, V-ATPase, mTOR, and MAPK/PI3K modules; nodes colored by first-appearance time point. Member-level data, Supplementary Data 7.

To identify the biological functions and androgen-sensitive members of the extranuclear proximal interactome, we performed functional enrichment analysis with g:Profiler across the cytosolic and microsomal compartments[58], uploading the shared proximal interactors at each time point in multi-query mode and querying Gene Ontology, KEGG, Reactome, WikiPathways, and CORUM (Figure 4C). We highlighted the categories with the largest temporal shifts in p-values to identify androgen-sensitive interactions with pathways previously implicated in AR signaling and pathways not previously implicated. These highlighted categories recapitulated well-established changes in chaperone-binding events that occur acutely before and immediately after androgens bind AR[60], and uncovered late-stage transcriptional effects that occur after AR undergoes cytoplasmic-nuclear translocation. For example, cytoskeletal protein binding was enriched at 0, 2, and 10 minutes and depleted at 30, 60, and 360 minutes, correlating with the known kinetics of AR cytoplasmic-nuclear trafficking[30], whereas the nucleocytoplasmic-transport pathway was depleted at 0 and 2 minutes and progressively enriched from 10 to 360 minutes, consistent with increased capture of transcription-associated interactions as liganded APEX2-AR translocated to the nucleus. The DNA damage response showed progressive enrichment at 30, 60, and 360 minutes[61,62,63,64], and the spliceosome and mRNA-processing categories increased from the 10-minute time point onward, demonstrating a progressive association with the splicing machinery involved in AR-dependent alternative splicing[20,21,65]. These findings establish AR-proximal interaction networks (AR-PINs) that are functionally linked to AR activity as a ligand-activated nuclear hormone receptor.

Beyond these well-characterized androgen-regulated processes, we examined smaller enrichment changes that might reveal novel AR-proximal interactions. Strikingly, the CORUM Retromer complex emerged as an androgen-sensitive group of proximal interactors whose enrichment decreased progressively with androgen exposure. The retromer complex mediates endosomal protein recycling to the trans-Golgi and plasma membrane[66,67] and has not been formally implicated in AR-dependent signaling. The WP Translation factors and the CORUM eIF3 complex similarly decreased at the 30-and up to the 360-minute time points, which is notable given that AR negatively regulates the expression of the protein-synthesis machinery[24].

To visualize the AR-PINs underlying these early trafficking enrichments, we constructed STRING physical interaction networks in Cytoscape from the shared extranuclear proximal interactors at each time point, filtered to the retromer, retrograde transport, and mTOR categories, and clustered by connectivity (Figure 4D). In the no-hydrogen-peroxide control, the network was dominated by ribosomal proteins, splicing factors, and metabolic enzymes with no detectable retromer or trafficking components, confirming that these networks are specific to APEX2-AR labeling rather than purification artifacts (Supplementary Fig. 4). At the earliest labeled time point (0 minutes), the network comprised a densely interconnected core of all retromer subunits (*i.e.*, VPS26A, VPS26B, VPS29, VPS35), multiple sorting nexins (*i.e.*, SNX1, SNX2, SNX3, SNX5, SNX6, SNX12), the retromer-associated GAP TBC1D5, the COG tethering complex, V-ATPase subunits, and mTOR-pathway components (i.e., MTOR, RPTOR, AKT1S1, TSC2, DEPDC5, MIOS, WDR59), together with the NSF-NAPA-NAPG fusion machinery (Figure 4D). The co-clustering of retromer and mTOR members is consistent with the endosomal co-localization of both complexes, where retromer mediates cargo recycling and the Ragulator-Rag complex recruits and activates mTORC1 [68]. These AR-PINs persisted for 2 and 10 minutes, then progressively contracted, and by 360 minutes had contracted to a minimal retromer core (i.e., VPS29, VPS26A, SNX3, SNX5, SNX6, SNX12) (Supplementary Fig. 4). The coincident loss of mTOR and V-ATPase proximity, tracking AR cytoplasmic-nuclear translocation, indicates that as liganded AR disengages from endosomal membranes, it loses the mTOR machinery while transiently retaining the retromer apparatus. This androgen-sensitive AR-retromer proximity nominated the retromer as a candidate regulator of AR, which we tested directly below.

### The nuclear AR proximal interactome resolves into ordered functional modules

To dissect the biological architecture of the nuclear AR-PIPs, we performed multi-query g:Profiler functional enrichment on the per-time-point nuclear AR-PIP gene lists against seven annotation sources (*i.e.*, GO BP, MF, and CC; KEGG; Reactome; WikiPathways; and CORUM) using a custom background of all 6,117 detected proteins[58], which returned 2,661 unique terms significant at an adjusted p of at most 0.05 in at least one time point (Supplementary Fig. 5). The resulting heatmap revealed a sharply biphasic temporal organization of the nuclear AR-PIPs. At the earliest time points (*i.e.*, 0 and 2 minutes), translation-initiation machinery dominated; this signature gave way to transcription, chromatin remodeling, mRNA processing, and the DNA damage response at later time points, thereby defining a translation-to-transcription handoff within the nucleus.

To gain deeper insight into the transcription coregulators and complexes comprising this time-resolved handoff, we generated Cytoscape STRING physical interaction networks for the nuclear AR-PIPs at each time point (Figure 5) [69,59]. Six biologically coherent AR-PINs emerged across the nuclear AR-PIP population, organized along the temporal trajectory from nuclear import through productive transcription and downstream RNA processing. The nuclear pore complex and karyopherin transport machinery populated the AR-proximal proteome earliest, capturing AR during nuclear import (Figure 5A), followed by the transcription and translation machinery (Figure 5B), chromatin modifiers spanning the NuRD complex, polycomb PRC1 and PRC2, the INO80 and SRCAP remodelers, the BAF and ncBAF SWI/SNF complexes, and the ATRX and DAXX heterochromatin axis (Figure 5C), the alternative-polyadenylation machinery (Figure 5D), the mRNA export and TREX complex (Figure 5E), and the DNA damage response (Figure 5F) [61,62,63,64].

**Figure 5.**
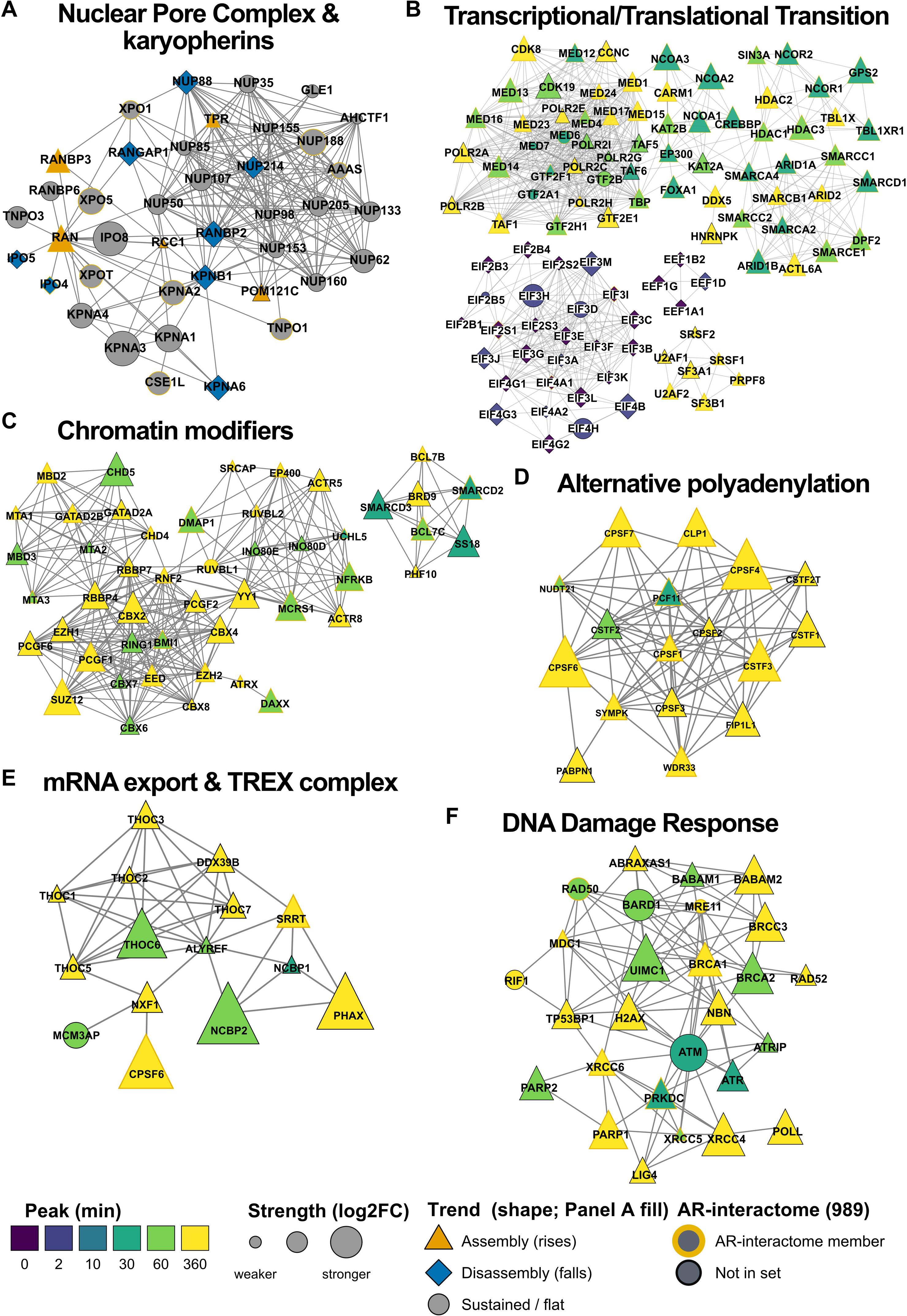
Cytoscape STRING physical networks of nuclear AR-proximal complexes across the androgen time course. (A) Nuclear pore complex and karyopherin transport (46 nodes), including NUP35-NUP214, AHCTF1, GLE1, AAAS, the RAN cycle, importins (KPNA1-6, KPNB1, IPO4/5/7/8, TNPO1/2/3), and exportins (XPO1-XPO7, XPOT). Captures AR during cytoplasmic-to-nuclear transit (peak 0-2 min). (B) Transcriptional/Translational Transition centerpiece, integrating the Mediator complex, RNA Pol II/GTFs, BAF and PBAF remodelers, p160 coactivators and corepressors, eIF3 and eIF4F translation complexes, and spliceosomal SR/U2 snRNP clusters. (C) Chromatin modifiers (48 nodes) including Polycomb PRC1, PRC2, NuRD, INO80/SRCAP, BAF/ncBAF, and the ATRX/DAXX axis. (D) Alternative polyadenylation (18 nodes): CPSF1-7, CSTF1-3, NUDT21, WDR33, FIP1L1, SYMPK, PCF11, CLP1, PABPN1. (E) mRNA export and TREX (15 nodes): THOC1-7, ALYREF, NXF1, NCBP1/2, MCM3AP, PHAX, DDX39B, CPSF6, SRRT. (F) DNA damage response (29 nodes): BRCA1/A complex, MRN, ATM/ATR/ATRIP, NHEJ, PARP1/2, BRCA2, RAD52, H2AX, TP53BP1, RIF1. Node shape and Panel A fill: Trend (triangle, assembly; diamond, disassembly; ellipse or circle, sustained). Node size: peak proximity log2FC (strength). Node color: Peak time point (min). Yellow border: 989-member AR-interactome membership. STRING physical subnetwork (see Methods for confidence cutoff and layout). Member-level data, Supplementary Data 9.

The temporal architecture resolved by these analyses recapitulated the canonical textbook model of nuclear receptor gene activation, in which ligand-bound AR nucleates ordered, cyclical waves of coactivator recruitment, chromatin-remodeler engagement, and RNA polymerase II loading at AR-regulated genes, with FOXA1 and ETS-family transcription factors serving as pioneer co-occupants at AR-bound enhancers across the AR cistromes [70,71,72,73,74,75,76]. Our nuclear PL-qMS time-course captured this canonical framework with high concordance, resolving Mediator, RNA polymerase II, and the pioneer factors within the nuclear AR-PIN and placing the canonical AR-at-enhancer axis on a much finer-grained map of chromatin-modifier deployment than previously reported.

### The retromer complex regulates AR-dependent transcription

The PL-qMS atlas established the retromer complex as AR-PIN of the broader AR-proximal interactome, with the three core subunits *(i.e.*, VPS35, VPS26A, VPS29) and associated sorting nexins called proximal across the androgen time course (Supplementary Fig. 6). To verify this proximity at endogenous protein levels without relying on APEX2-mediated biotinylation, we performed proximity ligation assays (PLA) targeting endogenous AR and the retromer subunits VPS26A, VPS29, and VPS35 in parental LNCaP cells using UnFold PLA probes (Figure 6A and 6B) [77], whose antibody conjugation was verified by silver-stained SDS-PAGE (Supplementary Fig. 7). Among the three AR-retromer pairings, the AR/VPS35 dual-probe condition produced the strongest PLA signal, with readily detectable foci at both the 0 and 60-minute time points, while all single-antibody controls exhibited minimal background, confirming that the dual-probe signals reflected true protein proximity rather than non-specific amplification. The strong AR/VPS35 signal is consistent with VPS35 serving as the retromer scaffold that contacts both VPS26A and VPS29 in the heterotrimeric core[78]. The PLA data thus confirms, at endogenous protein levels and in unmanipulated LNCaP cells, the robust, time-resolved AR-retromer proximity established by the atlas.

**Figure 6.**
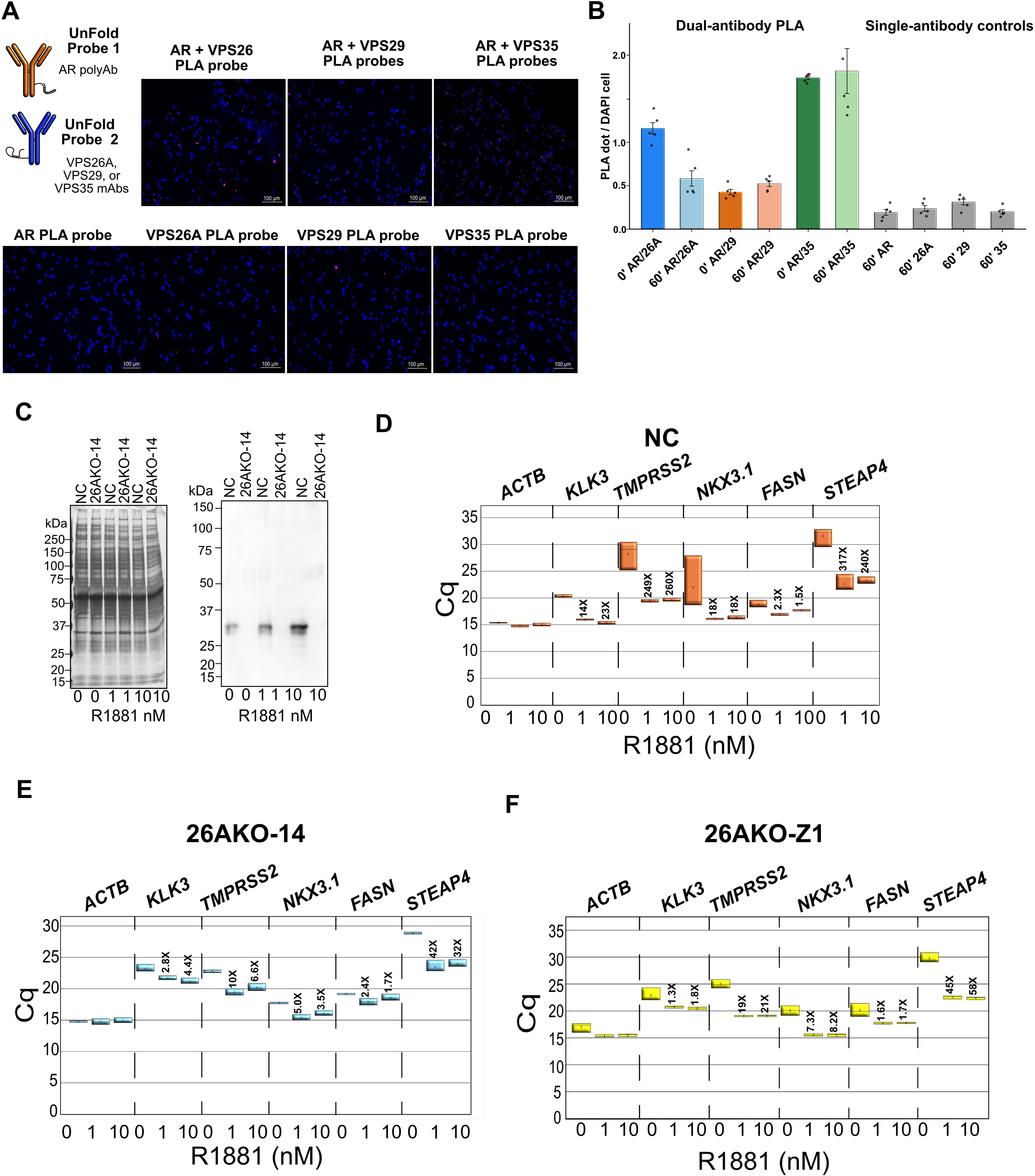
The retromer complex regulates AR-dependent transcription. (A) UnFold PLA probe schematic (Probe 1, AR polyclonal antibody; Probe 2, VPS26A, VPS29, or VPS35 monoclonal antibody) and representative PLA images in LNCaP cells. Top row, dual-antibody PLA (AR + VPS26, AR + VPS29, AR + VPS35); bottom row, single-antibody controls (AR, VPS26A, VPS29, VPS35). DAPI (blue), PLA Alexa-647 (pink); scale bars, 100 µm. (B) Quantification of PLA signals per DAPI-positive cell. Dual-antibody PLA (AR/VPS26A, AR/VPS29, AR/VPS35 at 0′ and 60′) compared with single-antibody controls (AR, VPS26A, VPS29, VPS35 at 60′); bars, mean ± standard deviation with individual measurements overlaid. (C) Silver stain and anti-KLK3 western blot of total cellular lysates from NC and 26AKO-14 cells across 0, 1, and 10 nM R1881. (D-F) Raw qPCR Cq values of androgen-regulated genes (ACTB, KLK3, TMPRSS2, NKX3.1, FASN, STEAP4) across 0, 1, and 10 nM R1881 in (D) NC, (E) 26AKO-14, and (F) 26AKO-Z1 cells; fold-change relative to vehicle is annotated per gene. PLA source data, Supplementary Data 10; blot and qPCR source data, Source Data file.

To test whether AR-retromer proximity had functional consequences, we developed the LNCaP-PB-luc2CP reporter cell line, in which the androgen-regulated rat probasin promoter drives luciferase, and confirmed dose-responsive androgen induction (Supplementary Fig. 8). Individual siRNA-mediated knockdown of VPS26A, VPS29, and VPS35 each reduced AR-dependent luciferase activity at both R1881 concentrations, with magnitudes comparable to co-knockdown of the classical AR coactivators NCOA1 and NCOA2, indicating that each core retromer subunit contributes to optimal AR-dependent transcriptional activity (Supplementary Fig. 8). Transfected siRNAs targeting the retromer subunits did not cause gross differences in cellular proliferation (Supplementary Fig. 9), indicating that the reduced AR-dependent luciferase activity did not reflect impaired cell growth. Because transient knockdown reduced retromer subunit levels only modestly (approximately 40 to 50%) and produced equivocal effects on endogenous KLK3/PSA (Supplementary Fig. 10), we sought orthogonal evidence, reasoning that if retromer integrity is required for AR transcriptional activity, then AR-driven prostate tumors should exhibit negative selective pressure against loss-of-function alterations in retromer genes. Querying cBioPortal across seven prostate cancer datasets encompassing 2,487 tumors, we found that deep deletions of VPS26A, VPS29, and VPS35 were essentially absent (0.1 to 1.3%) and point mutations were virtually absent, whereas the retromer-interacting but non-core sorting nexin SNX3 was deleted in 30 to 39%, demonstrating locus-specific purifying selection that paralleled selection at the AR locus itself (Supplementary Fig. 11).

The equivocal knockdown results prompted us to employ CRISPR-Cas9 genome editing for stable genetic disruption. We generated LNCaP cell lines harboring loss-of-function mutations in VPS26A and recovered two independent partial-knockout clones, 26AKO-14 and 26AKO-Z1 (Supplementary Fig. 12). Complete loss of VPS26A or VPS35 was not tolerated in LNCaP cells, consistent with the hypotetraploid LNCaP karyotype and essential retromer function [79]. Western blot analysis confirmed near-undetectable KLK3/PSA protein in the knockout cells despite largely unchanged VPS26A, VPS35, and AR protein levels, demonstrating that a gross reduction in retromer or receptor abundance did not underlie the transcriptional defect in KLK3/PSA (Figure 6C); densitometric quantification of Western blots used ECL detection validated within its linear range (Supplementary Fig. 13).

To assess AR-dependent transcription directly, we measured a panel of canonical androgen-regulated genes (*i.e.*, KLK3, TMPRSS2, NKX3.1, FASN, and STEAP4) by quantitative PCR in vehicle- and R1881-stimulated Negative Control (NC), 26AKO-14, and 26AKO-Z1 cells (Figure 6D-6F). In NC cells, androgen induced these genes to markedly different extents, most strongly TMPRSS2 and STEAP4 and only weakly FASN. In both Cas9-targeted VPS26A mutant clones, the androgen induction of every strongly responsive gene was substantially attenuated, whereas the weakly responsive FASN was largely unchanged and the ACTB control was stable across cell lines and treatments. These genetic experiments demonstrate that partial loss of VPS26A is sufficient to attenuate AR-dependent transcription of a broad panel of canonical androgen-regulated genes, indicating that intact retromer expression is required for optimal androgen-regulated gene expression in LNCaP prostate tumor cells.

### Partial VPS26A disruption mislocalizes the AR coactivator TMF1

To investigate why AR-dependent transcription was attenuated in the Cas9-targeted VPS26A mutant clones, we examined the localization of the membrane and organelle markers transferrin receptor (TFRC), early endosome antigen 1 (EEA1) (Supplementary Fig. 14), Golgi matrix protein 130 (GM130), and transcription factor modulatory 1 (TMF1) by immunofluorescence microscopy. Because the retromer complex recycles proteins from endosomes to the trans-Golgi network and cell surface, we focused on the Golgi markers GM130 and TMF1. GM130 and TMF1 displayed strong punctate Golgi co-localization in NC cells, but this co-localization was largely absent in both Cas9-targeted VPS26A clones, where the juxtanuclear punctate staining characteristic of intact Golgi localization was replaced by a diffuse cytoplasmic pattern (Supplementary Fig. 15).

The disrupted Golgi localization of TMF1 prompted us to assess whether its total abundance was also altered. Western blot analysis revealed no pronounced differences in TFRC, EEA1, TMF1, or AR protein levels across the three cell lines, indicating that the mislocalization phenotype reflects altered subcellular distribution rather than changes in protein abundance, and KLK3/PSA was again undetectable in both clones (Supplementary Fig. 16). Given that TMF1 is a coactivator of AR-mediated transcription[7] and our previous identification of a functional link among COPI retrograde trafficking, TMF1, and AR-mediated transcription[47,80], we examined AR-TMF1 co-localization across the androgen time course (Figure 7). In NC cells, AR displayed the expected cytoplasmic-to-nuclear translocation upon R1881 stimulation, and TMF1 showed progressive nuclear accumulation with intensifying AR/TMF1 co-staining at the 60 and 360-minute time points. In both Cas9-targeted VPS26A clones, AR nuclear translocation was intact, but TMF1 staining remained diffuse throughout the cytoplasm and the androgen-induced intensification of nuclear AR/TMF1 co-staining observed in NC cells was absent, paralleling the attenuated androgen-regulated genes measured in the same clones (Figure 6).

**Figure 7.**
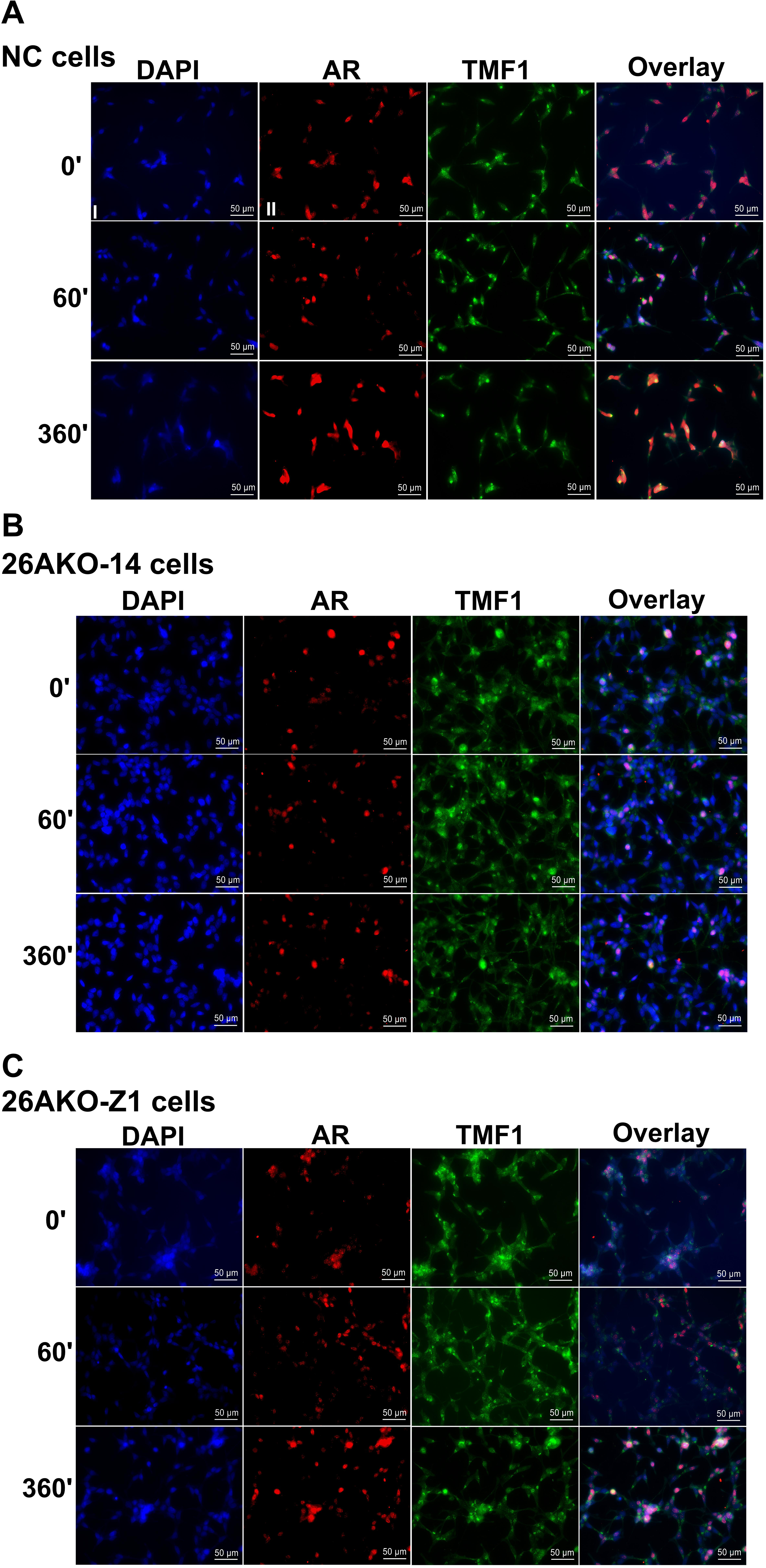
Mislocalization of TMF1 in VPS26A mutant cell lines. (A-C) Fluorescent microscopy for DAPI (blue), AR (red), and TMF1 (green), with overlay, in (A) NC, (B) 26AKO-14, and (C) 26AKO-Z1 cells after 48 hr propagation in androgen-depleted growth medium (10% charcoal-stripped fetal bovine serum) and challenge at 0′ with ethanol and at 60′ and 360′ with R1881 androgen. Scale bars, 50 µm.

Together, these results suggest a model in which the retromer complex, through VPS26A, maintains the Golgi localization of TMF1 and ensures its availability as an AR coactivator during androgen-stimulated transcription. We speculate that retromer disruption disperses the Golgi-confined TMF1 pool into a diffuse cytoplasmic distribution, impairing ligand-induced nuclear co-localization with AR. A failure to engage TMF1 in the nucleus is predicted to attenuate transcription of canonical androgen-regulated genes. This would establish the retromer complex as a functional modulator of AR-dependent gene expression and a specific extranuclear AR-PIN that shapes androgen-regulated transcription.

### Nuclear AR engages the translation machinery and validates against the AR chromatome

The nuclear networks revealed an early, transient engagement of AR with the cap-binding translation initiation machinery, a signature we validated orthogonally by PLA. Performing UnFold PLA in LNCaP cells[77], proximal interactions between AR and eIF4G were verified. eIF4G is a scaffolding subunit of the eIF4F cap-binding initiation complex, and 4E-BP1 is the mTORC1-regulated translational repressor that gates eIF4E availability[81,82]. Interestingly, both eIF4G and 4E-BP1 produced specific signals to AR, which were above single-antibody controls (Figure 8A). These findings confirmed the early nuclear engagement of AR with translation cofactors captured by the atlas.

**Figure 8.**
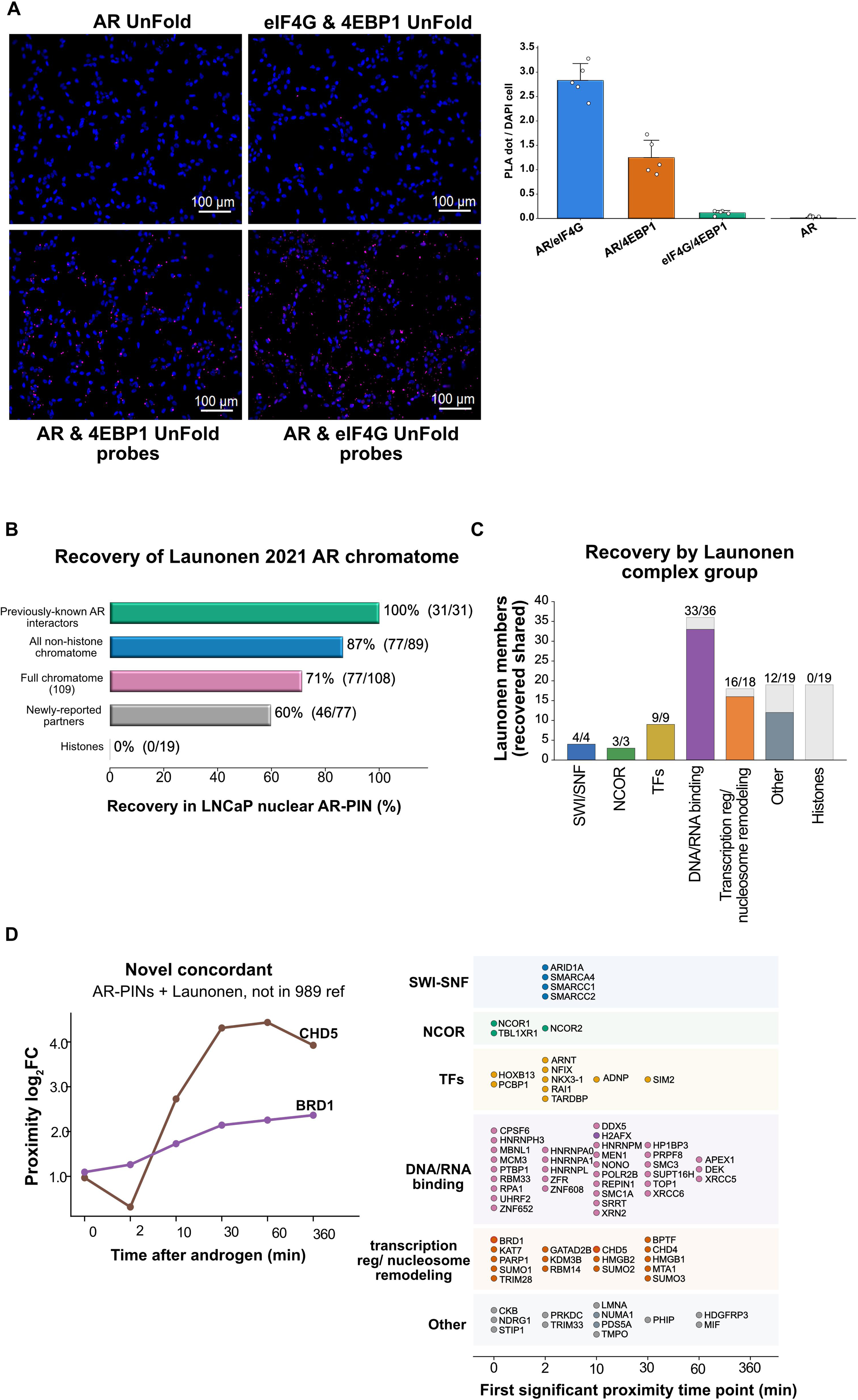
Nuclear AR engages the translation machinery and cross-validates against the AR chromatome. (A) Representative UnFold PLA images and quantification in androgen-depleted LNCaP cells at the 0-min time point. Images: AR alone (single-antibody control), eIF4G + 4E-BP1 (irrelevant-pair control), AR + 4E-BP1, and AR + eIF4G; DAPI (blue), PLA Alexa-647 (pink); scale bars, 100 µm. Bar graph, PLA dots per DAPI-positive cell (AR/eIF4G, AR/4E-BP1, eIF4G/4E-BP1, and AR); mean ± SD with individual values. (B) Recovery of the Launonen 2021 ChIP-SICAP chromatome in the LNCaP nuclear AR-PIN, stratified by subset. Previously known AR interactors, 100% (31/31); all non-histone chromatome, 87% (77/89); full chromatome, 71% (77/108); newly reported partners, 60% (46/77); histones, 0% (0/19). (C) Recovery by Launonen complex group. SWI/SNF (4/4), NCOR (3/3), transcription factors (9/9), DNA/RNA binding (33/36), transcription regulation and nucleosome remodeling (16/18), other (12/19), and histones (0/19); colored segment, recovered; gray, not recovered. (D) Left, novel concordant hits proximal in both datasets and absent from the 989-member AR-interactome reference (CHD5, BRD1), shown as log2 fold-change across 0-360 min. Right, temporal swimlane of the first significant proximity time point for each recovered Launonen-named complex member, colored by group as in (C). Member-level data, Supplementary Data 11; PLA source data, Source Data file.

### Orthogonal cross-validation recovers and extends the chromatin-engaged nuclear AR-PIN

To benchmark the nuclear AR-PIP catalog against an orthogonal, chromatin-directed proteomic method, we cross-validated it against the 109-protein AR chromatome reported by Launonen and colleagues, which was generated by ChIP-SICAP [44]. This biochemical technique combines chromatin immunoprecipitation with on-chromatin DNA biotinylation and streptavidin pull-down to isolate proteins co-bound to chromatin with androgen-bound AR in VCaP cells (Figure 8B). PL-qMS in LNCaP cells recovered the Launonen chromatome with high concordance across every stratification examined. All 31 previously validated AR interactors in the reference were recaptured (100%, 31 of 31), non-histone recovery was 87% (77 of 89), full-chromatome recovery was 71% (77 of 108), and 60% of the newly reported Launonen partners were recovered, whereas histones, which lack sequence-specific AR proximity, were not (Figure 8B). Recovery was uniformly high across all Launonen complex groups, including SWI/SNF (4 of 4), NCOR (3 of 3), transcription factors (9 of 9), DNA and RNA binding proteins (33 of 36), and transcription regulators and nucleosome remodelers (16 of 18) (Figure 8C).

We also extended the cross-validation beyond this network and recovered the chromatin remodelers CHD5 and BRD1, which were independently confirmed as AR-proximal in both datasets, despite their absence from the curated 989-member AR-interactome reference. Both remodelers increased in abundance over the androgen time course in our PL-qMS experiments (Figure 8D). These findings highlight two advances that distinguish the PL-qMS catalog from the static Launonen chromatome. Firstly, PL-qMS resolves the time of arrival of each recovered Launonen member across the androgen time course, a kinetic dimension that ChIP-SICAP cannot deliver (Figure 8D). Secondly, PL-qMS captures four families of nuclear cofactors entirely absent from the chromatin-directed dataset, namely translation-initiation machinery (n = 18, including 4E-BP1), mRNA processing and alternative-polyadenylation factors (n = 18), mRNA export and TREX (n = 15), and the nuclear pore complex and karyopherins (Supplementary Fig. 17). Together, these comparisons validate the chromatin-engaged nuclear AR-PINs as a reproducible biological feature. It also suggests that PL-qMS is an orthogonal method that extends the nuclear AR proximal proteome beyond the chromatin-resident reference set by ChIP-SICAP.

## Discussion

Proximity labeling captures both direct interactors and neighboring proteins of a target receptor and, when coupled with quantitative mass spectrometry, identifies hundreds to thousands of proximal interactors [12]. The field has often dismissed these neighbors as uninformative bystanders, reflecting general receptor metabolism rather than functional biology [39]. We argue the opposite. Two lines of evidence indicate that the breadth of the AR proximal interactome, across all three compartments, reflects biology rather than background. First, the curated AR-interactome is significantly over-represented within the proximal proteome at every time point and in every compartment (Figure 3), and the proximal landscape is reproducible across orthogonal proximity-labeling platforms, different cell lines, and labeling chemistries. Random contamination would neither enrich a curated interactome nor reproduce across independent methods. Second, the proximal interactome is best understood not as an expanded binary partner list but as cellular cartography, a functional neighborhood. APEX2 generates a labeling radius of approximately 200 nm [34], encompassing direct binding partners, transient interactors, and spatially proximal effectors whose organization around AR reflects a signaling architecture rather than random diffusion. Not every AR-PIP needs to engage AR functionally; a protein positioned within the AR proximal neighborhood nonetheless carries the latent potential to influence AR signaling should its expression or localization become dysregulated, as frequently occurs in human prostate cancers [13,30].

Why should AR maintain such a large functional neighborhood? Three converging features of AR biology make this expected rather than anomalous. AR is a hub protein with an extensive intrinsically disordered N-terminal domain, and disorder-rich hubs engage many low-affinity, transient, and often multivalent interactions that binary assays such as yeast two-hybrid and co-immunoprecipitation systematically miss, but that proximity labeling readily captures [27,83]. Liganded AR further partitions into biomolecular condensates [84,40], and condensate co-partitioning concentrates a large, dynamic protein population in the vicinity of AR, producing a proximal proteome that far exceeds the stoichiometric binary interactome. Finally, interactomes are expression-dependent and context-specific [85], so a deep, fractionated, time-resolved survey in androgen-responsive cells necessarily recovers more than any single-context binary screen. Together, these properties explain why the AR proximal interactome is large and continues to grow without reaching saturation, and why that breadth is a feature of hub-receptor biology rather than an artifact.

Two distinct modes of AR engagement contribute to this breadth. In the high-affinity, structured-domain mode, bona fide AR coregulators assemble on the AR N-terminal domain through structured interaction surfaces, with p160 and SRC-family coactivators engaging mapped non-LXXLL surfaces and Mediator-complex coactivators concentrated at AR N-terminal condensates [86,87,88]. In the transient, low-affinity short-linear-motif mode, a much larger cloud engages the AR AF-2 charge clamp competitively; the AR AF-2 groove is biochemically receptive to LXXLL motifs in vitro [89,90], and the human proteome contains thousands of LXXLL-bearing proteins. Each individual LXXLL engagement is brief and low-affinity, and the AR-specific FXXLF (FQNLF) motif of the AR N-terminal domain dominates AF-2 occupancy intramolecularly once the N/C interaction commits AR to chromatin engagement. This second mode is therefore permissive primarily before AR commits to its DNA-bound state, consistent with proximity labeling capturing both the curated AR coregulator core and a much larger short-linear-motif-reachable cloud sampled during cytoplasmic-to-nuclear transit. The proteome-wide LXXLL distribution across cytosolic, microsomal, and nuclear AR-PIPs, and its predicted divergence in the ligand-binding-domain-deleted variant AR-V7, are developed in a companion bioinformatic study [91].

Beyond cataloging, the atlas yields concrete mechanistic advances. In the extranuclear compartment, PL-qMS detected 82.2% of the known AR-interactome (813 of 989) and verified 81.9% of detected members as significantly enriched AR-PIPs, establishing the most comprehensive spatiotemporal map of extranuclear AR-proximal interactions to date. Retromer complex enrichment, including VPS29, RAB7A, and VPS35, emerged within 10 minutes of androgen stimulation and persisted throughout the time course; partial genetic disruption of VPS26A attenuated transcription of canonical androgen-regulated genes, demonstrating that a proximal interaction identified by PL-qMS has direct functional consequences. The mechanism converges on TMF1, an established N-terminal AR coactivator whose Golgi localization is disrupted in VPS26A-mutant cells [7], thereby linking retromer-mediated membrane trafficking to AR transcriptional output. The discovery of retromer exemplifies the utility of PL-qMS to identify functionally important AR-proximal interactions that traditional binary methods miss.

The identification of retromer subunits as AR-proximal proteins is consistent with growing evidence that VPS35 and associated factors function beyond their canonical role in endosomal cargo sorting [67]. VPS35 localizes to the nucleus, where it promotes transcription through interactions with transcriptional regulators at target promoters [92], and retromer-associated factors engage the Ku70/Ku80 heterodimer and DNA-PKcs at DNA double-strand break sites to facilitate non-homologous end joining [93,94]. The detection of VPS26A and VPS35 as AR-proximal proteins may therefore reflect not only endosomal encounters during AR trafficking but also nuclear co-residence, where retromer subunits could interface with AR-dependent transcriptional programs or DNA damage responses, a question that future work will resolve.

In the nucleus, the most unexpected finding is the transient, early-time-point engagement of AR with the cap-binding translational-initiation machinery, with direct AR-eIF4G and AR-4E-BP1 contacts verified by proximity ligation (Figure 8). This mechanistically extends the work of Liu and colleagues, who established that AR represses cap-dependent translation initiation in castration-resistant prostate cancer by transcriptionally upregulating 4E-BP1 [24]. Our proximity-ligation-verified contacts establish a parallel physical axis on the same proteins, with transient kinetics consistent with AR docking briefly on eIF4F upon nuclear arrival before releasing to engage chromatin. Loss of AR in castration-resistant disease would derail this coupled axis on both fronts, potentially explaining why AR-low tumors exhibit the most dramatic eIF4F hyperactivity and sensitivity to disruption of eIF4E and eIF4G [95]. More broadly, three non-traditional mechanisms beyond canonical enhancer-driven gene expression are represented in the nuclear-proximal AR proteome, namely alternative polyadenylation, alternative splicing, and cap-dependent translation control [13,23], thereby converting the predicted expanded repertoire of AR actions into a temporally resolved catalog of specific protein partners for each axis.

The nuclear proximal proteome also resolves chromatin-modifier engagement with unprecedented granularity, identifying 48 chromatin-modifying proteins distributed across six functionally distinct complexes spanning polycomb PRC1 and PRC2, NuRD, the INO80 and SRCAP remodelers, BAF and ncBAF SWI/SNF, and the ATRX and DAXX heterochromatin axis. Exactly half of these proteins are known members of the AR-interactome, providing internal validation, while the remainder represent previously uncatalogued proximal chromatin partners of AR. The simultaneous resolution of these complexes within a single proximity proteome places the canonical AR-at-enhancer transcriptional axis on a much finer-grained map of chromatin modifier deployment than previously reported [96].

Integrated across compartments and over time, these findings define a single, contiguous AR-proximal landscape (Figure 9). Androgen triggers AR engagement within the extranuclear retromer, trafficking, and mTOR machinery at the membrane and in the cytosol. AR then transits the perinuclear and nuclear pore compartments, where translational cofactors hand off to the nuclear machinery, and it finally assembles a transcription-competent complex in an ordered sequence from pioneer factors through chromatin remodelers to Mediator and RNA polymerase II. The translational cofactors are AR-proximal only transiently at the earliest time points and are lost as the transcription machinery assembles, marking the switch from a translation-cofactor-dominated to a transcription-dominated proximal interactome. The atlas thus renders AR signaling as one spatial and temporal continuum from the cytoplasmic membrane to chromatin rather than as isolated compartments, and the assembly of AR-PIPs into functional complexes at each stage constitutes AR-proximal interaction networks (AR-PINs), of which the retromer-AR-TMF1 axis and the AR-eIF4G-4E-BP1 axis are the two functionally validated examples reported here.

**Figure 9.**
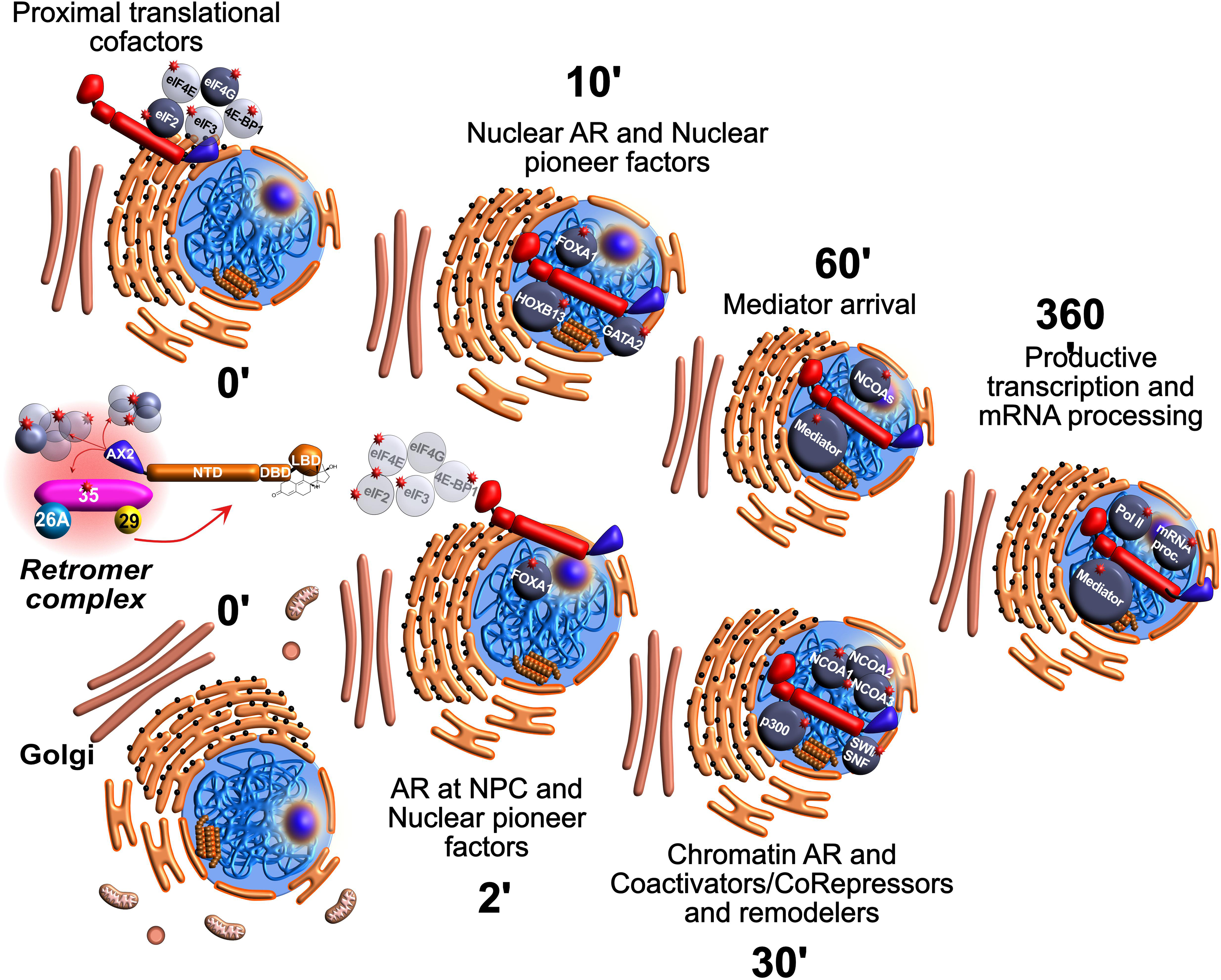
A unified spatiotemporal model of the AR proximal interactome. Integrated schematic of the AR proximal interactome across the androgen time course (0 to 360 min after R1881). At the plasma membrane, R1881 enters the cell and APEX2-AR (AX2) biotinylates proximal proteins (biotin tyramide; red stars mark biotinylated AR-PIPs). In the cytosol, AR engages the retromer complex (i.e., VPS35, VPS26A, VPS29) among the endosomal, lysosomal, proteasomal, Golgi, and mitochondrial compartments; AR domains (NTD, DBD, LBD) and the APEX2 (AX2) tag are indicated. At 0 min, proximal translational cofactors (i.e., eIF4E, eIF4G, eIF2, eIF3, 4E-BP1) are AR-proximal at the endoplasmic reticulum and nuclear periphery. AR then transits the nuclear pore complex and engages nuclear pioneer factors (FOXA1 at 2 min; FOXA1, HOXB13, and GATA2 at 10 min). By 30 min, chromatin-associated AR recruits coactivators, corepressors, and remodelers (i.e., p300, NCOA1, NCOA2, NCOA3, SWI/SNF). Mediator arrives by 60 min, and by 360 min AR assembles a transcription-competent complex driving productive transcription and mRNA processing (Mediator, RNA polymerase II, and mRNA-processing machinery).

The AR proximal landscape is reproducible across methods. Comparison with an independent Mini-TurboID-AR proximal proteome generated in PC3 cells converged on a shared core of 417 proximal interactors despite differences in cell line, labeling enzyme, and design [84] (Supplementary Fig. 18), and extension to BirA-AR datasets in LAPC4 and HEK293 cells recovered 203 of 268 and 25 of 39 AR-proximal proteins, respectively, within our extranuclear fractions [97,98]. The intersection across all four AR proximity datasets defines an 18-protein conserved core comprising the BAF and SWI/SNF chromatin-remodeling complex, canonical AR corepressors, and the H3K9 demethylase JMJD1C [99,100,101,102], captured by every AR proximity labeling method tested regardless of compartment, cell line, or enzyme (Supplementary Fig. 19). Cross-validation against the chromatin-directed Launonen AR chromatome recovered all 31 previously validated interactors (100%) and 86.5% of the non-histone chromatome [44], confirming the chromatin-engaged core of the nuclear AR-PIN as a reproducible biological feature, while the complementary spatial coverage of proximity labeling explains why our dataset uniquely captures the transient, non-DNA-binding translation machinery that chromatin-directed methods cannot see.

The temporally resolved AR-PIN catalog is therapeutically actionable. Recent AR-transactivation-domain inhibitors disrupt AR-coregulator partnerships including SWI/SNF, FOXA1, SMRT, and TBL1XR1 [103,104,25,105, 26], all of which populate our AR-PIN at the sampled time points, suggesting that the AR-PIN architecture defines the actionable surface of the AR transcriptional machinery and that pharmacological disruption of specific AR-PINs might selectively attenuate AR transcriptional programs in prostate cancer.

An important methodological consideration is whether the bait protein is expressed from the endogenous locus or from an inducible transgene. Endogenous tagging ensures physiological expression but means every AR molecule constitutively carries the bulky fusion tag, potentially driving clonal selection over successive passages. We employed doxycycline-inducible APEX2-AR, which captures a temporal snapshot of AR-proximal interactions without chronic adaptation to an engineered receptor and preserves the endogenous receptor, at the cost of a mixed population of tagged and untagged AR. The high recovery of the known AR-interactome and the strong cross-validation with orthogonal proximity-labeling datasets indicate that the proximal interactome captured here faithfully reflects physiological AR biology rather than overexpression-driven artifacts.

## Limitations of this study

Several limitations merit consideration. First, CRISPR-mediated disruption of VPS26A was incomplete owing to the hypotetraploid karyotype of LNCaP cells [79], and the extent to which partial VPS26A loss impairs retromer assembly remains undefined. Complete genetic ablation will be needed to determine the full extent of retromer-dependent AR transcriptional regulation. Second, the AR-proximal interactome was defined in a single androgen-sensitive prostate cancer cell line, and different androgen-responsive models will express distinct populations of AR-PIPs at varying abundances. Proximal partners that engage AR only in the context of ligand-binding domain-deleted splice variants or castration-resistant settings fall outside the immediate scope of this dataset and are natural targets for follow-up in AR-V7-expressing and castration-resistant models. Third, proximity ligation verifies physical proximity but does not establish direct binding or functional consequences; structure-guided point mutations of the AR AF-2 groove and N-terminal coactivator-binding surfaces will be needed to test the two-mode engagement model directly. Fourth, the temporal sampling resolves events on the minutes-to-hours scale, leaving faster nuclear-arrival events and longer-term remodeling outside our window. Beyond the proximity-ligation-verified and genetically validated axes reported here, the atlas is presented as a discovery resource for the field to interrogate, including the motif and domain grammar of the AR-PIP set and a compartment-resolved proximity interaction database developed in companion studies [91,106].

## Methods

Key resources, including antibodies, cell lines, chemicals, kits, and software, are provided in the Key Resources supplementary table.

### Cell Lines

LNCaP clone FGC (ATCC CRL-1740; RRID:CVCL_1379; authenticated by short tandem repeat (STR) profiling at the source) human prostate adenocarcinoma cells were maintained in RPMI 1640 medium supplemented with 10% fetal bovine serum (FBS), 1% penicillin-streptomycin, and 1% L-glutamine at 37°C in a humidified atmosphere containing 5% CO2. For androgen-depletion experiments, cells were propagated in phenol red-free RPMI 1640 supplemented with 10% charcoal-stripped fetal bovine serum (CSS) for 24–72 hours as specified for each experiment. The parental LNCaP clone was tested for mycoplasma contamination (InvivoGen) prior to cell line derivation. To generate LNCaP-APEX2-AR cells, the pTRE-Tight vector (Clontech) was first modified by replacing the low-copy origin of replication with a high-copy one, resulting in pTRE-Tight2 (pTRE-T2). The microRNA-adapted (mIR) cassette from pCDNA6.2-GW/miR (Invitrogen) was subcloned into the EcoRI–XbaI sites of pTRE-T2, and site-directed mutagenesis was performed to engineer unique BsaI sites for directional cloning of Block-iT microRNAs and to remove non-unique restriction sites (XhoI, XbaI, BsaI). Puromycin and hygromycin resistance cassettes were inserted via NheI–EagI sites using In-Fusion cloning, generating pTRE-T2-miR-PURO (Addgene #102646) and pTRE-T2-miR-HYGRO (Addgene #102647). To create the APEX2-AR expression construct, the mIR cassette was removed from pTRE-T2-miR-HYGRO and replaced with a multiple cloning site. The APEX2 coding sequence was obtained from pcDNA3-APEX2-NES (Addgene #49386) and a gene synthesis construct incorporating an N-terminal streptavidin-binding peptide (SBP) tag, a Kozak consensus sequence, and engineered restriction sites was appended to the APEX2 coding sequence. The N-terminal SBP-APEX2 cassette was subcloned into the modified pTRE-T2 hygromycin backbone, yielding NewNTERM_APEX2_HYGRO (5,343 bp). Full-length human AR cDNA was then amplified from pcDNA3-N-term-AR and subcloned into the AsiSI–MluI sites, to generate the pTRE_T2_deltamIR_NTERM_APEX2_AR_HYGRO (8,105 bp). This construct produces an N-terminal SBP-APEX2-AR fusion protein that retains all functional AR domains (NTD, DBD, hinge, and LBD). All constructs were verified by Sanger sequencing. Primer sequences used for PCR amplification and subcloning are listed in Supplementary Table 1. The final APEX2-AR expression construct will be deposited in Addgene. The parental LNCaP-Tet3G cell line (clone-19) was generated by transfecting parental LNCaP cells with the pCMV-Tet3G On neomycin expression vector (Clontech) and selecting for neomycin-resistant (e.g. 500 ug/ml) clones that demonstrated doxycycline-inducible expression using the pTRE3G-Luc reporter plasmid. LNCaP-Tet3G clone-19 cells were then transduced with the pTRE-T2-NTERM-APEX2-AR construct, selected with hygromycin (100 ug/ml), and stable clones were screened for doxycycline-inducible expression of APEX2-AR, leading to the discovery of LNCaP-APEX2-AR clone-13 cell line. To generate the LNCaP-PB-luc2CP polyclonal cell line, the proximal rat probasin (PB) promoter was PCR subcloned from rat genomic DNA into the EcoRI-XhoI multiple cloning site of the promoter-less expression vector pGL4.16[luc2CP]-Hygromycin containing the destabilized luciferase reporter gene (luc2CP). Parental LNCaP cells were transfected with the PB-luc2CP expression vector, selected for hygromycin resistance (150 ug/ml), pooled clones were expanded, tested for androgen-mediated luciferase induction, and subsequently designated LNCaP-PB-luc2CP cells. NC and VPS26A mutant cell lines (26AKO-14 and 26AKO-Z1) were generated from parental LNCaP cells by CRISPR-Cas9 genome editing as described below.

### Proximity Labeling and Subcellular Fractionation

LNCaP-APEX2-AR cells, grown in charcoal-stripped serum (CSS) medium for 72 hours, were treated with 10 ng/ml doxycycline added during the final 24 hours to induce APEX2-AR expression. Cells were incubated with 500 µM biotin-tyramide (biotin-phenol) for 1 hour at 37°C and treated with 100 nM synthetic androgen R1881 for defined time points (i.e., 0, 2, 10, 30, 60, and 360 minutes). Biotinylation was initiated by adding hydrogen peroxide (H2O2) to the growth medium to a final concentration of 1 mM for 60 seconds, alongside a negative control group of cells treated with PBS for 60 seconds. The biotinylation reaction was quenched by washing the cells three times with quench buffer (i.e., PBS supplemented with 10 mM sodium ascorbate, 5 mM Trolox, and 10 mM sodium azide). Cells were scraped off the 500 cm² tissue culture plates into chilled 50 ml conical tubes and then centrifuged at 2000 rpm at 4°C in a swing-bucket tabletop centrifuge for 10 minutes. The liquid was carefully decanted to avoid disturbing the pelleted cells, and the cell pellet was resuspended in hypotonic buffer (10 mM EPPS, 1.5 mM MgCl_2_, 10 mM KCl, pH 7.9, supplemented with 2 mM DTT and 1X HaltTM protease inhibitors), incubated on ice for 10 minutes, and then subjected to nitrogen cavitation at 100 PSI for 2 minutes. The lysate is centrifuged at 3,000 rpm for 20 minutes at 4°C. The supernatant, which contains the cytosolic and microsomal protein fractions, is collected and subjected to 100,000 × g ultracentrifugation for 3 hrs to pellet the crude microsomes, and the supernatant is collected as the cytosolic fraction. The nuclear pellet remaining after the 3000-rpm centrifugation step was collected and stored at -80°C. Cytosolic samples were desalted into a binding buffer (100 mM EPPS, 400 mM NaCl, pH 8.5) with Zeba desalting spin columns to remove residual small molecules and biotin-tyramide, and samples were quantified by the BCA method. Crude microsomal pellets were solubilized in a denaturing binding buffer (100 mM EPPS, 400 mM NaCl, 1% SDS, pH 8.5), and solubilized microsomes were quantified with the BCA method. For the nuclear fraction, the nuclear pellet retained after the 3,000 rpm centrifugation was extracted by the Dignam method[107]. Each nuclear pellet was resuspended in nuclear extraction buffer (NEB; 20 mM EPPS, 420 mM NaCl, 1.5 mM MgCl2, 25% glycerol, pH 7.9, supplemented with 2 mM DTT, 1x Halt protease inhibitors, and 750 U/ml Benzonase) at approximately 2 ml NEB per time-point pellet, incubated for 1 hour at 4 C on an end-over-end nutator to release chromatin-bound proteins, and centrifuged at 13,000 rpm in an SS-34 rotor (4 C, 25 minutes). The cleared nuclear extract was collected, diluted in denaturing binding buffer (100 mM EPPS, 400 mM NaCl, 4 M urea, 1% SDS, pH 8.5), and quantified by the BCA method.

### Streptavidin Affinity Purification

Equal amounts of input protein for each time-point (i.e., ∼5 mg) from both cytosolic and membrane preparations were diluted into denaturing binding buffer (100 mM EPPS, 400 mM NaCl, 4M Urea, 1% SDS, pH 8.5), samples were reduced with 5 mM TCEP for 15 minutes at room temperature, alkylated with 10 mM chloroacetamide for 25 minutes at room temperature in the dark, and quenched with 5 mM DTT for 15 minutes. Reduced and alkylated samples were incubated overnight at room temperature with pre-washed Pierce magnetic streptavidin beads (300 µl suspension volume; catalog 88817) in denaturing binding buffer. Beads were collected on a magnetic platform for 10 minutes, washed 5 times with 500 µl of denaturing incubation buffer. After the final wash, the beads were released from the magnetic platform, 300 µl of elution buffer (200 mM EPPS, 5% SDS, pH 8.5) was added to the sample, and the samples were incubated at 95°C for 5 minutes. The samples were incubated on the magnetic platform for 60 seconds to capture the magnetic beads, and a 1/10 portion was collected for Western blot analyses. The remaining portion of the sample was removed from the magnetic platform, allowed to resuspend to form a homogeneous bead sample, and subjected to single-pot solid-phase-enhanced sample preparation (SP3), adapted from the Proteomics Unit at the University of Bergen. Sera-Mag SpeedBeads were prepared by washing to remove sodium azide and reconstituted at a working concentration of 100 mg/ml. SP3 beads (300 µg) were added to the SDS-eluted sample at a 10:1 bead-to-protein ratio (wt/wt), and protein binding was induced by adding 100% acetonitrile to a final concentration of 80%, followed by incubation at room temperature for 10 minutes at 1,000 rpm. After centrifugation at 16,000 X g for 5 minutes and magnetic separation, beads were washed three times with 80% ethanol. On-bead digestion was performed overnight at 37°C in 100 µl of digestion buffer (200 mM EPPS, pH 8.5, 1 mM CaCl2) containing trypsin at a 1:25 enzyme-to-substrate ratio, with the addition of 10 pmoles of Pierce Retention Time Calibration Mixture heavy-labeled peptides as an internal standard. Digested peptides were recovered by magnetic separation; beads were subjected to a single wash in 500 mM NaCl. The collected samples were pooled, acidified, and desalted using Pierce peptide desalting spin columns (catalog 89852). Desalted tryptic peptides were fractionated by high-pH reversed-phase chromatography using Pierce C18 peptide desalting spin columns (catalog 89851). Dried peptides were resuspended in 300 µl of Buffer A (10 mM ammonium bicarbonate, 5% acetonitrile) and loaded onto pre-conditioned C18 columns. Unbound peptides were collected, and bound peptides were eluted in a stepwise acetonitrile gradient of 5%, 10%, 15%, 20%, 25%, 30%, 35%, 40%, 60%, and 90% acetonitrile in 10 mM ammonium bicarbonate, generating 11 fractions. Fractions were pooled non-contiguously to maximize peptide diversity. The unbound fraction and fractions 1, 4, 7, and 10 were pooled; fractions 2, 5, and 8 were pooled; and fractions 3, 6, and 9 were pooled, yielding three final pooled samples per biological replicate. Pooled fractions were dried by vacuum centrifugation and desalted using Pierce C18 spin tips (catalog 84850) prior to LC-MS/MS analysis.

### Mass Spectrometry

Peptides were analyzed by LC-MS/MS using a nanoElute 2 liquid chromatograph coupled to a timsTOF Pro2 mass spectrometer (Bruker Daltonics). Samples were loaded on a capillary C18 column (25 cm length, 150 µm inner diameter, 1.5 µm particle size, 120 Å pore size; PepSep Ultra). The flow rate was maintained at 500 nL/min. Solvent A was 0.1% formic acid in water, and Solvent B was 0.1% formic acid in acetonitrile. Peptides were separated on a 60-minute analytical gradient from 3% to 35% Solvent B for a total run time of 70 minutes. The timsTOF Pro2 was operated first in dda-PASEF mode to generate spectral libraries. MS and MS/MS spectra were acquired from 100–1700 m/z. The inverse reduced ion mobility (1/K0) was set to 0.70–1.40 V·s/cm2 over a ramp time of 100 ms. Data-dependent acquisition was performed using 10 PASEF MS/MS scans per cycle with a near-100% duty cycle. Ions were fragmented at an ion-mobility-dependent collision energy, which was linearly increased from 20 to 59 eV in positive ion mode. Low-abundance precursor ions with an intensity above a threshold of 500 counts but below a target value of 20,000 counts were repeatedly scheduled and otherwise dynamically excluded for 0.4 minutes. Each cytosolic biological replicate (n=3) was interrogated by dda-PASEF to generate a comprehensive cytosolic spectral library via MSFragger. A single microsomal biological replicate was used to generate a microsomal spectral library. The two spectral libraries were combined to generate a cytosolic-microsomal spectral library. dda-PASEF data were uploaded to py_diAID (https://github.com/MannLabs/pydiaid) to optimize scan windows for DIA acquisition. The timsTOF Pro2 was then operated in dia-PASEF mode[36] using 12–25 scan windows, based on py_diAID-optimized window parameters, for data-independent acquisition across all three biological replicate samples from both the cytosolic and microsomal protein fractions. The inverse reduced ion mobility (1/K0) was set to 0.70–1.40 V·s/cm2 over a ramp time of 100 ms with 100% duty cycle.

### Proteomic Data Analysis

Raw timsTOF Pro2 data were analyzed using FragPipe combined with MSFragger[51] and Philosopher[108] for dda-PASEF spectral library generation and DIA-NN for dia-PASEF protein quantification (FragPipe 24.0, MSFragger 4.4.1, DIA-NN 1.8.2 beta 8), applied identically to all three fractions[53,54,55,56]. dda-PASEF raw files were searched against a human UniProt FASTA database[109] (canonical sequences, no isoforms) supplemented with sequences for streptavidin, PRTCM, and an APEX2-AR fusion protein in which the APEX2 coding sequence was appended to the AR entry (UniProt P10275). A target-decoy database was generated for FDR estimation. MSFragger search parameters were set as follows, with all other settings left at default values. Enzyme was set to stricttrypsin (cleaving after K/R) with one missed cleavage permitted, peptide length 7–50 residues, peptide mass range 500–5,000 Da, precursor mass tolerance ±20 ppm with mass calibration and parameter optimization enabled, fragment mass tolerance 20 ppm, and isotope error 0/1/2. Variable modifications included oxidation of methionine (+15.9949 Da), N-terminal acetylation (+42.0106 Da), and phosphorylation of serine, threonine, and tyrosine (+79.9663 Da, max 3 per peptide). Cysteine carbamidomethylation (+57.0215 Da) was set as a fixed modification.

Peptide validation was performed with PeptideProphet. Spectral libraries were generated using Pierce iRT (Peptide Retention Time Calibration Mixture, containing heavy-lysine- and heavy-arginine-labeled peptides) as the retention-time calibration standard. DDA-derived spectral libraries were processed through py_diAID[52] (https://github.com/MannLabs/pydiaid) to optimize dia-PASEF scan windows, with a precursor m/z range of 200–1,450 and an ion mobility (1/K0) range of 0.70–1.40 V·s/cm², using 100 iteration steps and 20 starting points for window optimization. dia-PASEF data were quantified using DIA-NN (1.8.2 beta 8) within FragPipe 24.0, with variable modifications passed via command-line options. Protein quantification data were analyzed and visualized with a custom, version-controlled R statistical pipeline that implements established differential abundance methods, namely variance-stabilizing normalization[110] , limma moderated t-tests[111] , and Benjamini-Hochberg FDR correction. DIA-NN protein-group matrices were filtered to proteins with at least two valid values in at least one condition, VSN-normalized, and missing values imputed by a down-shift model (width 0.3, downshift 1.8 SD, seed 20260601) with each injection treated as a technical replicate. Differential proximal enrichment was tested per timepoint versus the −H₂O₂ control with limma moderated t-tests and Benjamini–Hochberg FDR correction. Proximal AR-PIP, adjusted P ≤ 0.05 and log2 fold-change > 1. The pipeline reads the raw DIA-NN matrix directly; no third-party statistical-GUI outputs were used. The nuclear fraction was processed through the identical FragPipe 24.0 and DIA-NN workflow, with protein-level intensities taken from the report.pg_matrix.tsv output. As a nuclear-specific acquisition detail, each nuclear biological replicate (R1 to R3) was acquired with three technical injections per time point; the R3 batch was initially acquired at reduced timsTOF Pro2 sensitivity and re-injected after instrument cleaning, and superseded injections were discarded (22 of 85 nuclear injections dropped). Each retained injection was treated as an independent technical replicate, yielding nine observations per time point, and nuclear protein groups were retained if they had at least two non-missing values in at least one time point (6,117 of 6,164 protein groups).

### Hypergeometric Enrichment Testing

Hypergeometric tests assessed the over-representation of the curated AR-interactome among proximal AR-PIPs. The AR-interactome reference was assembled from BioGRID, NCBI, and published interaction databases[3], yielding 1,010 unique gene symbols; 24 obsolete HGNC symbols (for example GNB2L1 to RACK1, KIAA1429 to VIRMA, WHSC1 to NSD2, and H2AFY to MACROH2A1) were resolved to current symbols via org.Hs.eg.db (Bioconductor 3.18), and after de-duplication the final reference contained 989 unique current-HGNC symbols. One-sided hypergeometric tests were applied at each time point, separately for the cytosolic, microsomal, and nuclear fractions, comparing this 989-member reference against the set of proteins enriched as proximal AR-PIPs (adjusted P <= 0.05 and log2 fold-change > 1).

### UpSet Plot Analysis

UpSet plot analysis was performed to characterize the temporal architecture[57] of the overlapping cytosolic-microsomal proximal AR-interactome across all six time points (0, 2, 10, 30, 60, and 360 minutes).

### cBioPortal Genomic Analysis

Genomic alteration data were retrieved from cBioPortal for Cancer Genomics across seven prostate cancer datasets encompassing 2,487 tumors spanning primary/localized disease and metastatic castration-resistant prostate cancer (mCRPC). Copy number alteration frequencies and mutual exclusivity analyses were performed for VPS26A, VPS29, VPS35, AR, and SNX3.

### Functional Enrichment Analysis

Gene-set functional enrichment of the androgen-responsive proximal interactomes was performed with g:Profiler (g:GOSt) through the gprofiler2 R package[58] (v0.2.4). For each androgen-stimulation timepoint (0, 2, 10, 30, 60, and 360 min, each compared against the no-H2O2 background control), the set of significantly enriched proximal interactors (AR-PIPs; Benjamini–Hochberg-adjusted P ≤ 0.05 and log2 fold-change > 1, deduplicated to current HGNC gene symbols) was submitted as an ordered query. Over-representation was tested against the default Homo sapiens genome-wide annotated gene universe (no custom experiment-specific background). Multiple-testing correction used the g:SCS (Set Counts and Sizes) algorithm, the native g:Profiler method, at a significance threshold of 0.05. The annotation sources queried were Gene Ontology (Biological Process, Molecular Function, Cellular Component), KEGG, Reactome, WikiPathways, and CORUM. The cytosolic∩microsomal overlap interactome was analyzed identically. Gene symbols throughout the pipeline were normalized to current HGNC nomenclature using the org.Hs.eg.db Bioconductor annotation package (v3.18.0; Bioconductor 3.18). All differential expression and enrichment analyses were performed in R 4.3.2; a fixed random seed (set.seed(20260601)) was applied during down-shift imputation upstream of the limma differential test to ensure full reproducibility of the proximal-interactor inputs to enrichment.

### Cytoscape Network Analysis

Protein-protein interaction (PPI) networks were constructed using Cytoscape (v3.10.4)[69,59] with the STRING physical interaction database. For each time point, the gene symbols of the shared proximal interactors from the overlapping cytosolic–microsomal AR-PIPs (Supplementary Data 7) were entered as a list into the Multiple Proteins search field of the STRING web server (https://string-db.org), with organism set to Homo sapiens (taxon 9606). Physical interactions were retrieved using the following parameters, namely a network type of physical subnetwork, minimum required interaction score of 0.700 (high confidence), smart delimiters enabled, and a 1% false discovery rate (FDR) stringency applied on the functional enrichment side. The resulting STRING interaction network was exported via the Exports tab as a tab-separated values (TSV) file and imported into Cytoscape (v3.10.4) via File → Import Network from File. The resulting STRING enrichment table was filtered to retain six functional annotation categories, namely GO Biological Process, GO Cellular Component, GO Molecular Function, KEGG Pathways, Reactome Pathways, and WikiPathways. Redundant terms were removed using the STRING redundancy filter with a cutoff of 0.70. Networks were clustered using the Markov Cluster Algorithm (MCL) within the STRING plugin, with functional enrichment retrieved for each cluster using the genome as background. Singleton nodes (proteins with no physical interactions above the confidence threshold) were removed from the visualization. Clustered networks were arranged using the yFiles Organic Layout algorithm, followed by the yFiles Remove Overlaps function to improve readability. To highlight specific enriched pathways (e.g., retromer complex, mTOR signaling), the corresponding rows were selected in the PPI enrichment output table, which highlighted the associated gene nodes in the network. Edges between selected nodes were then selected (Select → Edges → Edges between selected nodes), and unselected nodes and edges were hidden to isolate the pathway of interest. This process was repeated for each enriched pathway visualized in the manuscript figures. Network images were exported as PNG files at 600 dpi resolution.

### Proximity Ligation Assay (PLA)

UnFold PLA probes were generated using SoluLink chemistry (Vector Labs Protein-Oligonucleotide Conjugation Kit, S-9011-1) adapted from the protocol described by Klaesson et al.[77]. Genscript synthesized Probe 1 TMO1-DNA oligonucleotide 5′-AMINE-AAAAATATGACAGAACTAGACACTCUUUCTATTAGCGTCCAGTGAATGCGAGTCCGTCTGAAAGAGTGTCTAGTTCTGTCATATTTAAGCGTCTTAA-3′ was conjugated to anti-VPS26A (Invitrogen, MA5-34664), anti-VPS35 (Santa Cruz, sc-374372), or anti-VPS29 (Santa Cruz, sc-398874), and Genscript synthesized Probe 2 TMO2-DNA oligonucleotide 5′-AMINE-AAAAAAGACGCTAATAGTTAAGACGCTTUUUAAAAAAAAGCGUCUUAACUAUUAGC GUC-3’ was conjugated to anti-AR polyclonal antibody (Proteintech, 22089-1-AP). Unfold probe antibody conjugation was verified by silver-stained SDS-PAGE (Supplementary Fig. 7), and 1/100 dilutions of Unfold Probe 1 and Unfold Probe 2 were used in PLA experiments. 1/50 dilutions of the single Unfold Probes were used in the negative control experiments. Approximately 15K LNCaP cells were seeded into each well of an 18-well chamber slide containing androgen-depleted medium (10% CSS) and grown for 48 hours. Cells were challenged with vehicle (i.e., ethanol) at 0’ or 10 nM R1881 synthetic androgen for 60’ and 360’. Cells were fixed in 10% buffered formalin, permeabilized with PBS containing 0.1% Triton X-100 for 10 minutes, and subjected to antigen retrieval in 100 mM Tris, pH 9.5, containing 6M urea at 80°C for 10 minutes. Slides were cooled at room temperature, washed twice for 5 minutes in Wash Buffer A (TBS-Tween 20, 10 mM Tris, 150 mM NaCl, 0.05% Tween-20, pH 7.4) and blocked for 1 hour at 37°C in a humidity chamber with in-house PLA blocking buffer (10% goat serum in PBS containing 0.1% ovalbumin, 0.02% sodium azide, and 100 µg/ml sheared single-stranded herring sperm DNA). The sheared ssDNA was prepared by diluting herring sperm DNA stock to 1 mg/ml, boiling at 95°C for 10 minutes, and snap-cooling on ice-ethanol to promote denaturation. After blocking, slides were washed three times for 3 minutes each in TBS-T (TBS with 0.05% Tween-20).

Primary antibody-DNA probes (TMO1 and TMO2) were diluted 1:100 to 1:300 in blocking buffer supplemented with 2.5 µg/ml sheared herring sperm ssDNA and incubated overnight at 4°C. Slides were washed three times for 3 minutes each in TBS-T, and the unfolding reaction was performed according to the modified protocol of Klaesson et al.[77]. PLA signals were quantified using BZ-X800 cell count software. The same UnFold PLA approach was applied to the nuclear AR and translational-cofactor pairs, with Probe 1 conjugated to anti-eIF4G and anti-4E-BP1 antibodies and Probe 2 conjugated to the anti-AR antibody.

### siRNA Transfection

siRNA transfections were performed using Lipofectamine RNAiMAX (Invitrogen) for 24-well format or Oligofectamine (Invitrogen) for 96-well format. For 24-well format, 47,000 cells per well were seeded in 500 µl of growth medium. 0.75 µl of 40 µM siRNA per well (30 pmoles; 50 nM final concentration) was diluted in 25 µl Opti-MEM, and 1.5 µl RNAiMAX per well was separately diluted in 6 µl Opti-MEM. The two mixtures were combined, incubated at room temperature for 5 minutes, diluted with 16 µl Opti-MEM, and 49 µl of the transfection complex was added per well. For 96-well format, 0.15 µl of 40 µM siRNA per well was diluted in Opti-MEM, and 0.3 µl Oligofectamine per well was separately diluted in Opti-MEM. The two mixtures were combined and incubated at room temperature for 25 minutes to allow complex formation, then diluted with additional Opti-MEM and added to cells (∼9.4 µl per well). LNCaP or LNCaP-PB-luc2CP cells were transfected with siRNAs at 50 nM (single target) or 25 nM each for co-targeting (NCOA1 and NCOA2). For luciferase reporter experiments, LNCaP-PB-luc2CP cells were androgen-starved for 24 hours, transfected with siRNAs for 48 hours, and restimulated with 10 nM or 100 nM R1881 for 24 hours prior to luciferase quantification.

### Luciferase Reporter Assay

AR transcriptional activity was measured using the LNCaP-PB-luc2CP reporter cell line. Dose-response was confirmed using 10-fold escalating concentrations of R1881 (0.001–1,000 nM). Luciferase activity was measured 24 hours after androgen stimulation.

### CRISPR-Cas9 Genome Editing

CRISPR-Cas9 genome editing was used to generate LNCaP cell lines harboring loss-of-function mutations in VPS26A. An sgRNA and a custom TrueTag Knockout Enrichment dsDNA targeting template (2,155 bp) introducing a premature stop codon within the VPS26A coding region were designed using the Invitrogen TrueDesign Genome Editor. The donor carried a neomycin resistance cassette for selection and a CMV-driven mRFP1 marker for identification of integrant clones. LNCaP cells (250,000 per well in 24-well format) were transfected using Lipofectamine CRISPRMAX. Tube 1 contained 1 µg TrueCut Cas9 Protein V2, 200 ng sgRNA, 250–400 ng dsDNA template resuspended in 10 µl Opti-MEM, 2.5 µl Cas9 Plus reagent, and 0.5 µl Alt-R HDR Enhancer V2 (1 µM working concentration). Tube 2 contained 1.5 µl Lipofectamine CRISPRMAX diluted in 25 µl Opti-MEM and incubated for 1 minute at room temperature. Tube 2 was added to Tube 1, mixed by pipetting, and incubated for 10–15 minutes at room temperature before adding ∼60 µl of the transfection complex per well of adherent cells. Drug-resistant clones were expanded, and on-target integration and the predicted premature stop codon at Ser120 were confirmed by genomic PCR and by Sanger sequencing of the subcloned integration junction. The donor cassette and the integrated-allele sequence are provided in Supplementary Data 15.

### Western Blotting

Protein lysates were resolved by SDS-PAGE and transferred to membranes. Membranes were probed with primary antibodies as specified in Supplementary Table 2 (Key Resources). Protein loading was verified by silver staining. Western blot signals were quantified by ImageJ densitometry. Linearity of ECL detection was validated (Supplementary Fig. 13).

### Immunofluorescence Microscopy

Cells were propagated in androgen-depleted medium (10% CSS) for 48 hours and challenged with 10 nM R1881 or ethanol vehicle. Cells were fixed in 10% formalin for at least 1 hour and stored in PBS at 4°C until staining. Slides were equilibrated to room temperature, washed in PBS containing 0.1% Triton X-100 for 10 minutes, and subjected to antigen retrieval by incubation in freshly prepared 100 mM Tris pH 9.5 containing 6M urea at 80°C for 10 minutes in a polypropylene Coplin container. After cooling to room temperature, slides were blocked with 10% FBS in PBS containing 0.1% ovalbumin for 1 hour. To validate doxycycline-induced APEX2-AR expression and proximity biotinylation, proximity-labeled cells were stained across the androgen time course for endogenous AR (AR441 mouse monoclonal antibody), APEX2-AR (anti-SBP mouse monoclonal antibody), and biotinylated proteins (streptavidin-HRP conjugate), with minus-H2O2 cells serving as the background-biotinylation control (Supplementary Fig. 1). For retromer and organelle marker immunofluorescence, secondary antibodies included goat anti-mouse Alexa Fluor 568 and goat anti-rabbit Alexa Fluor 488. After three PBS washes (5 minutes each), nuclei were counterstained by briefly dipping slides in 60 ml dH2O containing 5 µl of 5 mg/ml DAPI, followed by a brief rinse in dH2O. Slides were mounted with ProLong Diamond Antifade mounting medium and cured overnight in the dark. Fluorescent images were acquired on the Keyence BZ-X800E platform.

### Quantitative RT-PCR

Total RNA was extracted using the Qiagen RNeasy Mini Kit according to the manufacturer’s instructions. First-strand cDNA was synthesized from purified RNA using the SuperScript IV First-Strand Synthesis System (Invitrogen). Briefly, template RNA was combined with oligo(dT)20 primers and 10 mM dNTP mix, denatured at 65°C for 5 minutes, and chilled on ice. The RT reaction mix (5× SSIV Buffer, 100 mM DTT, Ribonuclease Inhibitor, and SuperScript IV Reverse Transcriptase at 200 U/µl) was added, and the combined mixture was incubated at 50–55°C for 10 minutes followed by heat inactivation at 80°C for 10 minutes. Quantitative PCR was performed using Qiagen RT² qPCR gene-specific primers for KLK3, TMPRSS2, NKX3.1, and FASN, custom primers designed against STEAP4 mRNA (NM_024636.4; see Supplementary Table 1), and ACTB as the reference gene. Delta-Ct fold changes were calculated by normalization to ACTB. Three independent biological replicate experiments were performed.

### Cell Proliferation Assay

LNCaP cells were transfected with siRNAs (50 nM). At 72 hours post-transfection, cellular proliferation was measured with the CyQUANT proliferation kit (Supplementary Fig. 9).

### Statistical analysis

Statistical analyses of proteomics data were performed with a custom R statistical pipeline. One-sided hypergeometric tests assessed AR-interactome enrichment, and differential abundance analysis used two-sided limma moderated t-tests with a Benjamini-Hochberg correction. PLA signals were quantified using BZ-X800 cell count software. Western blot signals were quantified by ImageJ. qPCR data represents three independent biological replicates with delta-Ct normalization to ACTB. Computational scripts for cross-validation overlap calculations were generated with AI assistance (Claude, Anthropic) and independently validated by the authors. All statistical tests and sample sizes are specified in figure legends.

## Data availability

The mass spectrometry proteomics data generated in this study have been deposited to the ProteomeXchange Consortium via the PRIDE partner repository[112] with the dataset identifier PXD081449. Upon publication, the dataset will be released to public access. Source data for the figure panels are provided in the Source Data file, and the full underlying datasets are provided in Supplementary Data 1 to 15.

## Code availability

Analysis code for the custom R statistical pipeline (variance-stabilizing normalization, limma moderated t-test, and Benjamini-Hochberg FDR correction) is deposited at https://github.com/mewrightlab/DIA-Toolkit. The repository is private during peer review, and reviewer access is available from the corresponding author. Upon manuscript acceptance, the repository will be made publicly accessible, a release tag (v1.0.0-AR-PINs) will be cut, and an archival Zenodo DOI ([Zenodo DOI pending]) will be minted and added to the published record. The pipeline was developed and tested on R 4.3.2.

## Materials availability

The LNCaP-APEX2-AR, LNCaP-PB-luc2CP, and VPS26A mutant (26AKO-14 and 26AKO-Z1) cell lines generated in this study are available from the corresponding author upon request. Plasmids, UnFold PLA probes, and other reagents generated for this study are available from the corresponding author upon completion of a Materials Transfer Agreement. The UnFold PLA antibody-DNA conjugates include the anti-AR, anti-VPS26A, anti-VPS35, anti-VPS29, anti-eIF4G, and anti-4E-BP1 probes.

## Acknowledgements

We thank Cathrine Pfukwa for assistance with early evaluation of light microscopy data and experimental data analysis. We thank Steven P. Gygi and Joao A. Paulo for their contributions to the initial evaluation of TMT chemistries for the analysis of experimental samples. We also thank Alexey I. Nesvizhskii and Fengchao Yu for their early assistance with MSFragger and the potential use of FragPipeAnalyst during the initial stages of this research. This work was supported by the National Institutes of Health (R01GM143399).

## Author contributions

M.E.W. conceived and designed the study, developed the LNCaP-APEX2-AR cell line, the APEX2-AR vector, and all additional cell lines and reagents, performed all cell biology experiments, including proximity labeling, subcellular fractionation (cytosolic, microsomal, and nuclear), proximity ligation assays, CRISPR knockout generation, Western blotting, immunofluorescence microscopy, qPCR, and luciferase reporter assays, generated all MS samples, analyzed and interpreted data, and wrote the manuscript with input from all authors. C.C.P. managed liquid chromatography conditions and acquired all LC-MS/MS data on the timsTOF Pro2 platform. C.O. generated data visualizations and analysis code for quantification of proximity ligation assay data and prepared figure displays. J.K.E. contributed to the computational analysis of the mass spectrometry data and provided computational pipeline support for proteomic data analysis. L.R. provided access to the timsTOF Pro2 mass spectrometer, contributed expertise in SP3 sample preparation, and advised on methods for optimizing peptide identification and quantification. L.R. and M.E.W. co-supervised the work, contributed to data interpretation, and edited the manuscript. M.E.W. secured funding. All authors reviewed and approved the final manuscript.

## Use of generative AI

During the preparation of this work, the authors used Claude (Anthropic) to assist with manuscript drafting, editing, formatting, and reference management, and, under the authors’ supervision, to execute computational scripts for cross-validation overlap statistics (hypergeometric tests) using standard statistical methods and publicly available databases (BioGRID). All computational code and outputs were independently reviewed and validated by the authors against the primary experimental data. The authors reviewed and edited all AI-assisted content as needed and take full responsibility for the publication’s content.

